# Ancestral state reconstruction with discrete characters using deep learning

**DOI:** 10.64898/2026.03.19.712918

**Authors:** Anna A. Nagel, Michael J. Landis

**Affiliations:** One Brookings Drive, Department of Biology, Washington University in St. Louis, St. Louis, MO 63130, USA

**Author notes:** Corresponding author details here.

**Keywords:** phylogenetics, deep learning, statistical models, ancestral state reconstruction, discrete characters

## Abstract

Ancestral state reconstruction is a classical problem of broad relevance in phylogenetics. Likelihood-based methods for reconstructing ancestral states under discrete character models, such as Markov models, have proven extremely useful, but only work so long as the assumed model yields a tractable likelihood function. Unfortunately, extending a simple but tractable phylogenetic model to possess new, but biologically realistic, properties often results in an intractable likelihood, preventing its use in standard modeling tasks, including ancestral state reconstruction. The rapid advancement of deep learning offers a potential alternative to likelihood-based inference of ancestral states, particularly for models with intractable likelihoods. In this study, we modify the phylogenetic deep learning software phyddle to conduct ancestral state reconstruction. We evaluate phyddle’s performance under various methodological and modeling conditions, while comparing to Bayesian inference when possible. For simple models and small trees, its performance resembles the performance of Bayesian inference, but worsens as tree size increases. While phyddle still performs adequately for more complex models, such as speciation and extinction models, the estimates differ more from Bayesian inference in comparison with simpler models. Lastly, we use phyddle to infer ancestral states for two empirical datasets, one of the ancestral ranges of a subclade of the genus *Liolaemus* and ancestral locations for sequences from the 2014 Sierra Leone Ebola virus disease outbreak.

## Introduction

Biologists interpret phylogenetic patterns of ancestral states to test hypotheses concerning trait evolution and lineage divergence. For example, ancestral states are routinely used to investigate historical scenarios involving adaptive radiation (Schluter et al., 1997), ecological convergence (Mahler et al., 2013), biogeographical dispersal (Ree et al., 2005), and molecular evolution (Yang et al., 1995). Under ideal conditions, the actual ancestral states for particular characters of now-extinct species can be extracted with little interpretation through the fossil record (e.g. mollusk shells, mammal teeth), whereas fossils are rare or non-existent for many other lineages and characters (e.g. angiosperm floral traits, virus protein structures, bird songs). Ancestral state reconstruction (ASR) methods help fill this gap by generating the ancestral state(s) of a character or trait along a phylogeny, particularly for those ancestral taxa represented by internal nodes. From the phylogenetic modeling perspective, ASR is simply an estimation problem, where the objective is to obtain the probability of every state for every node, given the observed (tip) states of sampled taxa and the phylogenetic tree, while also assuming the estimation model is correct. For these reasons, our ability to convincingly test evolutionary hypotheses using ASRs is largely predicated upon the biological realism of our phylogenetic models, the quality of our data, and the statistical accuracy of our reconstruction methods.

With this in mind, a large body of work has developed statistical methods and corresponding software to estimate ancestral states (reviewed in Joy et al., 2016). Many popular phylogenetics programs can infer ancestral states of discrete characters, using DNA substitution models (Yang, 2007; Kozlov et al., 2019; Minh et al., 2020), models of historical biogeography (Ree and Smith, 2008; Matzke, 2014), or morphological evolution (Revell, 2012; Beaulieu et al., 2012; Pagel et al., 2004). The R packages DIVERSITREE and CASTOR (FitzJohn, 2012; Louca and Doebeli, 2018) can infer ancestral states for more complex models, such as state-dependent speciation and extinction (SSE) models (e.g. Maddison et al., 2007; Goldberg et al., 2011; Goldberg and Igić, 2012; Beaulieu and O’Meara, 2016). Many of these methods use empirical Bayes approaches to ASR, which first find the maximum likelihood estimates of the parameters, and then infer the ancestral states conditional on the exact parameter estimates (Yang et al., 1995). Other programs take a fully Bayesian approach and estimate the posterior distribution of ancestral states corresponding with samples from the posterior distribution of parameters (Höhna et al., 2016; Suchard et al., 2018; Bouckaert et al., 2019).

Key to standard ASR methods is the ability to derive and calculate the likelihood for the model of character evolution. For simple models of character evolution, such as Markov models, Felsenstein (1981) provided an exact and tractable method to compute the model likelihood, while for more complex models, such as state-dependent speciation-extinction (SSE) models, likelihoods must instead be computed numerically (Maddison et al., 2007). However, there exists a third class models that realistically describe biologically important processes, but models of this kind are not accompanied by tractable methods for computing likelihoods. For example, SIR models incorporate changing infection dynamics overtime (Kermack and McKendrick, 1927). Such SIR models and their derivatives do not have simple phylogenetic likelihood functions, though they can be approximated in some cases, such as during the initial phase of the outbreak (Featherstone et al., 2022) with birth-death models. Using coalescent theory (Volz et al., 2009), researchers have developed accurate approximations to the SIR model for reconstructing disease spread among locations beyond the initial outbreak phase (Müller et al., 2018, 2019). However, such innovations reflect the fact that, when compared to simpler models, complex models require extraordinary amounts of effort and insight to be rendered useful for application. Novel methods are required to estimate ancestral states for other models that have no known likelihood functions, for any number of theoretical or practical reasons.

Deep learning may offer a general solution to infer ancestral states for such intractable models, without requiring a likelihood function. Recent advances have addressed several key obstacles to using deep learning with trees, such as how to structure phylogenetic data into tensors so it may be used to train neural networks for classical modeling tasks (Voznica et al., 2022; Lambert et al., 2023). Additionally, new software such as phyddle has improved the accessibility of phylogenetic deep learning methods for users interested in likelihood-free modeling tasks (Landis and Thompson, 2025). More complete reviews of recent developments in deep learning in phylogenetics can be found elsewhere (Braichenko et al., 2025; Mo et al., 2024; Borowiec et al., 2022).

Advances such as these have laid the foundation to develop general ASR strategies using deep learning. ASR can be viewed as a classification task, with the goal of accurately predicting the exact state(s) for one or more internal node(s) in the tree (output) from the phylogenetic tree and tip-state patterns (input). Taking this perspective, Thompson et al. (2024) used supervised learning to train a neural network to predict the ancestral state of the root node, obtaining estimates as accurate as Bayesian inference under a variety of simulated epidemiological scenarios. This strategy treated the ancestral state for the root node as a standalone categorical variable, with no specific relationship to the tree topology; every tree in the training set, regardless of its size or topology, will have such a node. Even for trees that share a given number of leaf nodes, the internal nodes for one tree topology will not be intrinsically comparable to the internal nodes for a different topology. In this case, any ancestral state variable for an internal node being estimated will not, in general, represent the same phylogenetic position across different training examples. When tree sizes vary with respect to numbers of represented taxa, the sheer number of prediction targets (internal nodes) also varies among trees. Because supervised learning aims to maximize predictive accuracy of the network across all training examples, it may not be easy to adapt the naive strategy from Thompson et al. (2024) for ASR corresponding to other internal nodes, besides the deepest root and/or origin node.

Imagine training a neural network to estimate ancestral states using a dataset containing two of the three possible rooted, unlabeled topologies for a 5-tip tree – i.e., that includes only many examples for tree topologies A and B, but *not* for topology C, from Figure 1. Given a large enough training dataset, the neural network will likely be able to estimate the ancestral states for trees with topologies A and B reasonably well. However, it is unclear how the network will perform with trees of topology C. Only some nodes in tree C are analogous to nodes in trees A and B. For example, the subtree descendant from node 2 in tree C and node 2 in tree B have the same topology. In this case, a neural network may also be able to infer ancestral states for node 2 and its descendant, node 1, in tree C reasonably well. However, there are no such analogous nodes in trees A or B for node 3 in tree C, so the neural network may perform worse for this node. The topology of the entire tree impacts ASR, at least in a likelihood-based framework, but this toy example illustrates how local patterns may be similar even if the entire topologies are not identical. In the case of 5-tip trees, it would be trivial to simulate all three possible topologies, but the issue is exacerbated when considering that most training sets have larger trees and both random tree sizes and topologies. Together, this means that few of all possible trees are perfectly represented, even in the largest of training sets. Dense representation over phylogenetic patterns of branch length and tip-state variation may also be needed to accurately infer ancestral states for topologies not seen during training. Thus, a key challenge in using deep learning for ancestral state estimation is generating sufficiently diverse training data and providing it in a format that is conducive to learning patterns across trees.

**Fig. 1.**
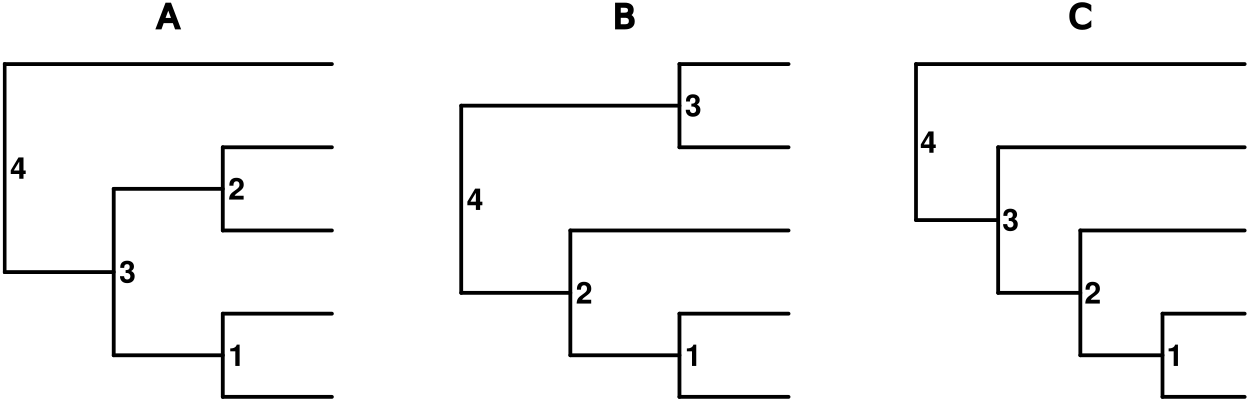
Trees A - C show the three possible unlabeled rooted topologies for a 5-tip tree. Each of the trees has four internal nodes. For each internal node, the goal is to estimate the ancestral state.

In this study, we develop new deep learning methods to estimate ancestral states of discrete characters and implement the methods in the program phyddle. We evaluate the performance of the methods for a variety of models – including Markov models, SSE models, and an SIR with migration model, which does not have a known likelihood function – and compare these results to Bayesian inferences when possible. To demonstrate how the method works under empirical settings, we use it to reconstruct the historical biogeography of *Liolaemus* lizards (Esquerre et al., 2019), and the ancestral spread of Ebola virus during the 2014 outbreak in Sierra Leone (Müller et al., 2019). In the discussion, we remark on the strengths and weaknesses of the current approach, and suggest how it may be improved in the future.

## Methods

Consider a rooted binary tree with known tip states for a discrete character. Our goal is to estimate the ancestral state of the discrete character at every node in the phylogeny for generic models of characters evolution using deep learning. We use the phyddle software, which was developed for deep learning with phylogenetic trees (Landis and Thompson, 2025). First, we describe phyddle and our approach to ancestral state estimation, including modifications to phyddle. Then, we describe the simulation methods to test the performance of phyddle for ASR. This included simulation experiments with four-tip trees and larger trees with a variety of models, including binary Markov, BiSSE, and GeoSSE models. To evaluate methodological performance, we compared the inferences against the true histories and against Bayesian estimates of the ancestral states. We assume Bayesian estimates represent peak attainable accuracy, using them as a reference to measure performance gaps for our deep learning approach. We compared both the most probable ancestral states and the probabilities of the ancestral states. These experiments should be taken as a rigorously documented baseline for ASR performance using a simple deep learning method. We conclude by analyzing two empirical datasets, demonstrating the flexibility of the method while illustrating where it performs well versus poorly.

### *Ancestral State Estimation with* Phyddle

Phyddle is a deep learning analysis pipeline for phylogenetics. It allows for inferences of discrete or categorical variables or parameters of any phylogenetic model that can be simulated. First, phyddle simulates input data using software of the users’ choice. Then, phyddle formats the data into tensors appropriate for machine learning. This includes encoding the phylogeny, tip states, parameters to be treated as data, and parameters to be inferred. Additionally, phyddle calculates a panel of summary statistics from the input data which are treated as data. Then, phyddle trains a neural network to estimate the parameters of interest. Finally, phyddle estimates parameters for the test dataset as well as any empirical data and plots the results. Landis and Thompson (2025) describes the design of phyddle in greater detail.

Phyddle was modified to allow for the estimation of ancestral states. phyddle uses compact bijective ladderized vector (CBLV) encoding for trees with serial-sampling or extinct tips (Voznica et al., 2022) and compact diversity-reordered vector (CDV) encoding for extant only trees (Lambert et al., 2023). These formats rotate the descendants of nodes on phylogenies by either the sample ages or branch lengths to encode the phylogenies into tensors. This rotation reduces the number of patterns the neural network needs to learn. Alternative formatting options in phyddle rotate the nodes similarly, but encode additional, redundant information in the tree tensors. The tip states are encoded with a modified CBLV+S or CDV+S format (Thompson et al., 2024). With this format, zero-padding allows for variable tree sizes up to a maximum using a fixed tensor size. To encode the internal node states as variables to estimate, the internal nodes are indexed following an in-order tree traversal on the formatted tree. The ancestral state at each node index is then treated as a categorical variable. For a tree with *N* tips, there are *N* − 1 internal nodes and thus *N* −1 categorical variables to estimate. The estimates for the ancestral states are mapped back to the original unformatted tree using internal node names provided in the input tree. Similar to the CBLV or CDV format, zero-padding is used for internal nodes that exist only in larger trees (Fig. S1).

As mentioned earlier, ASR for discrete characters can be viewed as a classification problem. Consider a model of character evolution with *S* states, where states cannot change at speciation events (e.g. simple Markov models). The objective is to assign probabilities to the *S* possible states for one or more of the *N* −1 internal nodes. To do so, we train our neural network through supervised learning using a cross-entropy loss function

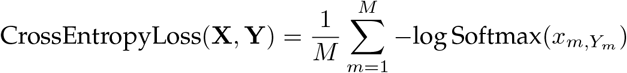

where **X** is a matrix of size *M* × *S* of support signals with values −∞ < *x*_*m,i*_ < +∞ for every state *i, Y* is a vector of labeled examples, such that *Y*_*m*_ ∈ {1, …, *S*}, and *m* indexes each of the *M* training examples. Our softmax function is defined as

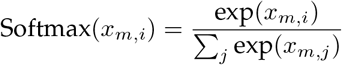

which converts *x*_*m,i*_, the signal supporting state *i* for example *m*, into a probability, such that 0 < Softmax(*x*_*m,i*_) < 1 and ∑_*i*_ Softmax(*x*_*m,i*_) = 1. As a consequence, the cross-entropy loss score contributed by example *m* improves when the network assigns a high probability that Softmax 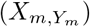 for the true state, *Y*_*m*_. This basic approach can be modified to target multiple nodes and more complex state spaces, as we describe below.

Using this general strategy, we consider three alternative ways to estimate ancestral states with phyddle. In the *marginal estimation* strategy, the ancestral state of each of the *N −*1 internal nodes is classified as a variable with *S* states, where each node has a Softmax function to report the probability of each state. For example, for a 4-tip symmetric tree with a binary character, the probability of state 0 and 1 would be estimated for each of the 3 nodes. This strategy is termed ‘marginal’ since each variable only reports ancestral state probabilities for its corresponding internal node, rather than (‘jointly’) for all nodes in the tree. Unless otherwise noted, the marginal strategy is used. Since phyddle estimates the same number of variables for each phylogeny, this means that if the maximum tree size for a network is *K*, there will be *K −*1 ancestral state estimates. The states for nodes that do not exist in the tree are zero-padded; zero-padded values are omitted when measuring accuracy. In the *joint estimation* strategy, a single variable with *S*^(*N*−1)^ states is estimated for each tree, using a single Softmax function. This corresponds to enumerating all possible combinations of internal node states on the tree. For example, for a 4-tip symmetric tree with a binary character, the probability of each of the 2^3^ possible combinations of internal node states would be estimated. To get the marginal probability of state 0 at the root node, for example, the probability of each state with 0 at the root are added. This corresponds to adding the probabilities of the trees with ancestral states {0,0}, {0,1}, {1,0}, and {1,1} at the non-root internal nodes. This strategy will suffer as the number of tips or states grows, since the number of categories grows rapidly. In the *single node estimation* strategy, the network is given the name of a single node for which to perform ASR. In this case, the network performs classification with one Softmax function for one node with *S* possible states. To estimate the probabilities of the ancestral states for all *N* −1 nodes, the estimation is run for each node independently. For example, to estimate the root state for a 4-tip symmetric tree with a binary character, the root is specified using a user provided node name, and the probabilities of states 0 and 1 are reported. Only one node per tree is used in the training dataset, so larger training dataset sizes may be required for the single node strategy to perform comparably to the marginal strategy.

For some models, such as GeoSSE (Goldberg et al., 2011), the character state can change at time of speciation. This leads to a triple of states for each internal node that correspond to the parental lineage before speciation, and the left and right daughter lineages after speciation. In principle, each of those states could be estimated independently using any of the previous strategies, using one network for the parent, one network for the left daughter, and another for the right daughter. However, we found this naive approach did not work well in practice. When trying to estimate the parental state, the accuracy of the inferences was consistently low, and the network seemed to be biased towards the most common states in the training data (results not shown). Alternatively, in the triplet strategy, all three states are estimated for each of the *N −*1 nodes, similar to the marginal estimation strategy (Fig. 2). For example, in Fig. 2a, before speciation at node 1, the ancestral lineage has the widespread range of A+B, and produces daughter lineages with ranges A+B and A. This cladogenetic state pattern for (A+B→A+B, A) would ultimately be encoded with the value of 6 (Fig. 2b-d). Ancestral state triplets are then predicted using Softmax functions, following the marginal strategy described above.

**Fig. 2.**
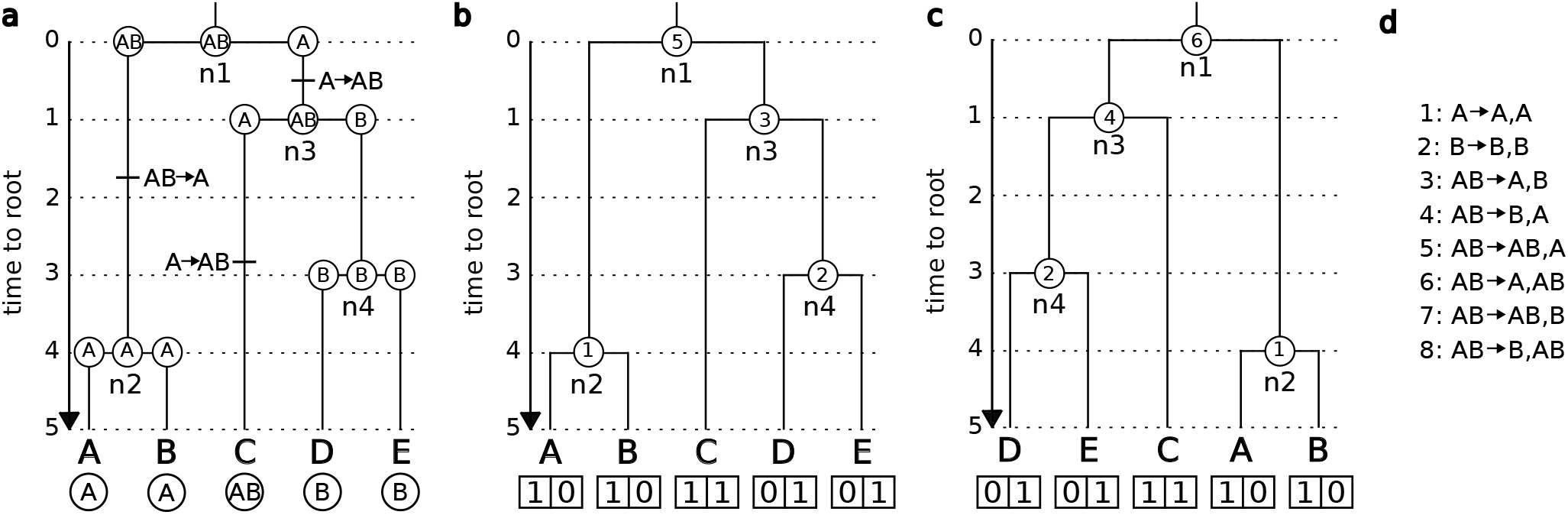
Tensor format for a phylogeny where states change at cladogenesis. (a) The biogeographic history is shown for a GeoSSE model. States can change along a branch or at the time of speciation. The history at each ancestral node is given by the states of the parent and both daughters. (b) Each event type at cladogenesis is coded as a number (d) and used to label the ancestral states in the tree. The labels differentiate between left and right daughters. The tip states are labeled as presence/absence in region A and B as two separate variables. (c) When the tree is formatted, some of the internal nodes in the tree are rotated. If the left and right daughters have different states, this changes the ancestral state. (d) The mapping for each possible state at an internal node in a tree in a GeoSSE model to a numeric value, written as parent→ left daughter, right daughter.

### Markov model of binary character evolution

#### Four-tip trees

Four-tip trees were simulated using a birth-death process (Nee et al., 1994) using the R package CASTOR (Louca and Doebeli, 2018). Log base-10 birth-rates were drawn from unif(−2, 0). To obtain the death rate, a rate was drawn from the same distribution and multiplied by the birth rate. This ensured the birth rate was higher than the death rate. The stopping condition was five taxa. If there were not five taxa, the simulation was rerun with new parameters. One of the two tips descendant from the last speciation event was dropped. This was to avoid zero branch lengths that result from ending the simulation with exactly four tips. Character data was simulated with a Markov model with two states with symmetric transition rates. Root states were drawn from the stationary distribution. Log base 10 transition rates were drawn from unif(−3, 0).

All three ancestral state estimation strategies were used (marginal, joint, and single node). The single node training dataset was three times larger to account for a smaller amount of training data per dataset (one vs three nodes). The first 50,000 trees were the same in all three datasets, and the single node dataset had an additional 100,000 trees. Each network was trained twice with the same dataset, which included using the same division between test and train datasets. Unless otherwise noted, the results reflect the inferences from one training. Estimates were compared for the same test datasets for all strategies. To directly compare the same test dataset for all three strategies, the test dataset from the marginal and joint strategies (which were the same) were treated as empirical data for the single node strategy. These trees were not in the training dataset for the single node network.

Bayesian inference of ancestral states for trees in the test dataset was performed using RevBayes (Höhna et al., 2016). The prior distributions for the transition, birth, and death rates were identical to those used for simulation. Two MCMCs were run for each analysis. MCMCs were run with a burnin of 2,500 iterations with a tuning interval of 100. After the burnin, the MCMCs were run for 25,000 iterations. The move assigned to each rate had weight of 2, meaning that two updates were proposed for the rate per iteration, on average. Convergence was assessed with ESS values and the Gelman diagnostic calculated using the R package CODA (Gelman and Rubin, 1992; Plummer et al., 2006). All but one pair of MCMCs had ESS values above 200 for the rate parameter in both MCMCs. All but one pair had a Gelman diagnostic below 1.1. The two pairs of MCMCs that did not meet this criteria were manually inspected in TRACER (Rambaut et al., 2018) and were considered to have converged based on very similar posterior distributions for the two runs.

To evaluate the concordance of the inferences from Bayesian inference and deep learning, the probabilities for each state as reported by Bayesian inference versus the three phyddle estimation strategies were compared. For comparisons based on point estimates, the most probable state was used as the point estimate. The point estimates for each method were compared to both the true ancestral states and the states inferred by the other methods.

To compare inferences among methods for particular topologies and internal node states, a larger dataset was generated with an additional 25,000 trees. Not all combinations of internal node states occurred with equal frequency, so the larger dataset gave greater representation to the rarer combinations. There are two topologies corresponding to rooted four-tip trees and eight combinations of internal node states for each of those topologies, making 16 internal node state-topology combinations in total. In a Markov model of character evolution with equal transition rates, the 0 and 1 labels for the character are arbitrary; state 0 could have been assigned to be state 1 and state 1 could have been state 0. For each of the sets of internal node states, there is a corresponding set of internal nodes where the 0 and 1 state assignments have been flipped (e.g. Fig. 3a trees 1 and 2). For a likelihood-based method, both of these setups should give equivalent results. However, it is possible this is not true for phyddle. To investigate this, for each tree in this dataset, the 0 and 1 labels were reversed to give a combined dataset with 50,000 trees. Inference of ancestral states was performed with phyddle using the previously trained networks. Since trees with the 0 and 1 labels reversed were included in the dataset, differences in performance across the reversed data would be due to biases of the method, not sampling error from the test dataset.

**Fig. 3.**
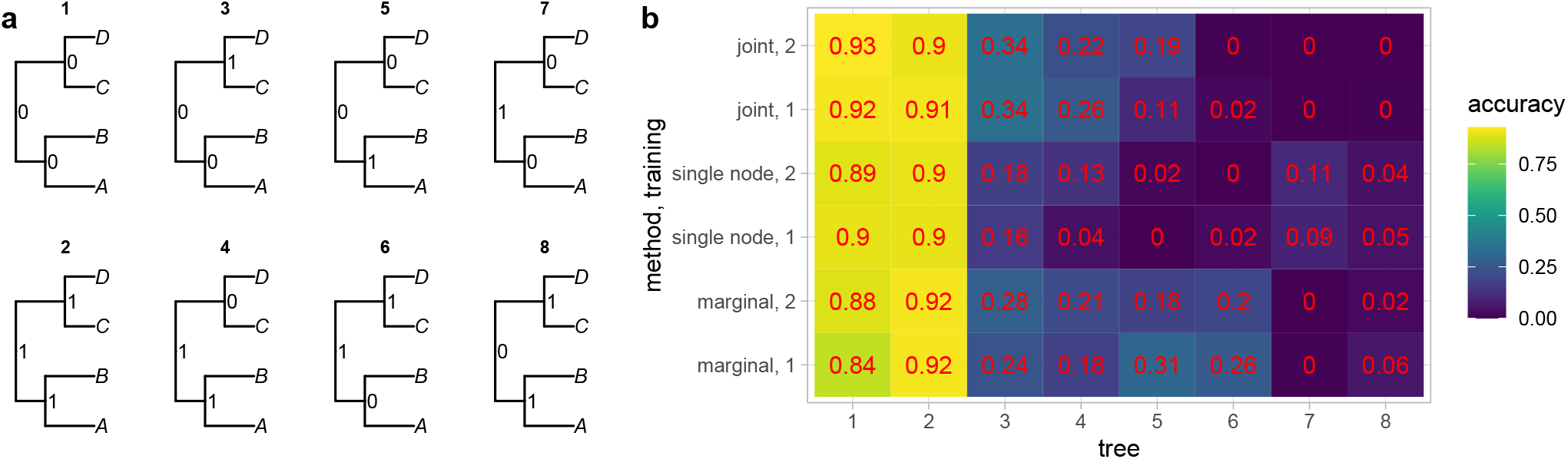
(a) There are 8 possible internal node state patterns for a symmetric four-tip tree with a binary character, shown in trees 1-8. The state patterns for the trees in the same column are symmetric; the only difference is the 0 and 1 labels have been switched on the internal nodes. When the trees are formatted with phyddle, the older daughter node is always the left daughter. In the first column, no changes are required on the internal branches to observe the ancestral state pattern. In the second column, at least one change is required to observe the pattern of ancestral states, and the change is on the longer internal branch. The third column also requires at least one change, but on the shorter branch. The last column requires at least two changes, one on each internal branch. (b) The heat map shows the proportion of inferences where all three ancestral nodes were correctly estimated for the corresponding tree and state pattern. The trees in the test dataset are classified into each of the 8 patterns of internal node states shown in (a) based on their true history. The x-axis shows the tree number, which matches the tree labels in (a), e.g. tree 1 has all internal nodes in state 0. Three strategies were used for inference, joint, single node, and marginal. In the joint strategy, a single categorical variable was estimated with possible states 1-8, corresponding the the 8 possible internal node state patterns in (a). For the single node strategy, the node name was given and the ancestral state for that node was estimated as a binary categorical variable. This was repeated three times to estimate the states for all nodes in the tree. In the marginal strategy, three binary categorical variables were estimated, each corresponding to one of the internal nodes in the tree. For each strategy, the neural network was trained twice. The numbers on each square match the values shown with the heat map colors.

#### Fixed tree size

Phyddle was used to simulate phylogenies and infer ancestral states for larger trees across three datasets with trees with different fixed numbers of tips. Phylogenies were simulated using the same method as for the four-tip trees but with the number of taxa for the stopping condition as 51, 101, or 201 tips to generate 50, 100, or 200 tip trees. The log transition rates between states were lower than the four-tip trees, drawn between unif(−3, −1), to reduce the chance of character saturation with larger (older) trees. Ancestral state estimation was performed with PHYDDLE using the marginal estimation strategy, training a separate network for each tree size. Additionally, ancestral state estimation for the root state alone was performed as in Thompson et al. (2024) using a separate network for each tree size.

Ancestral states were reconstructed with RevBayes. The priors were set to the same distributions as used for simulation. MCMCs were run for 25,000 iterations. The burnin was an additional 2,500 iterations and has a tuning interval of 100. Each rate was assigned a move with a weight of 2. Two MCMCs were run for each analysis. ESS values and the Gelman diagnostic were calculated with CODA. Datasets with pairs of MCMCs that did not have a Gelman diagnostic of less than 1.1 or where either MCMC did not have an ESS value of 200 were removed from the analysis. This resulted in removing no more than 59 of 2500 datasets for each tree size.

To investigate the accuracy of the probabilities reported by phyddle for the ancestral states, all of the internal nodes in the test datasets were binned by probability of state 1 in intervals of 0.1 ([0, 0.1), [0.1, 0.2) … [0.9, 1.0]). The average probability reported by PHYDDLE was calculated for each bin and compared to the proportion of nodes where state 1 was the correct state for each bin. To study performance based on the nodes’ positions in the tree, each node was binned by its height relative to the total tree height from [0, 0.1), [0.1, 0.2) … [0.9, 1.0]. The proportion of inferences where the most probable state was the true state was calculated for each bin for both the Bayesian and phyddle estimates.

#### Variable tree size

Three sets of simulations and inferences were made using phyddle with trees that were 10 to 50 tips, 20 to 100 tips, and 40 to 200 tips. The simulation design was the same as for the fixed tree size experiments, except the tree size was drawn uniformly between the minimum and maximum tree size for the dataset. If there was full extinction, all parameters and the tree size were redrawn. Ancestral states were estimated with RevBayes using the same set up as was used for the fixed tree size. Convergence was assessed in the same way as for the fixed tree size analyses. No more than 4% of each dataset was removed due to lack of convergence. The proportion of inferences where the Bayesian and phyddle inferences were correct across tree-height bins and the proportion of inferred states that were correct by the inferred probability bins were calculated as described above.

Different node numbers correspond to different (average) positions within the tree, where some nodes may be easier to infer than others. For instance, owing to the zero-padded region of the CDV format, the nodes with indexes greater than the number of internal nodes in the smallest trees in the training dataset always have less training data than lower node numbers for the variable size trees, which may impact the accuracy of inferences. The proportion of inferences where the point estimate was correct for the test dataset was compared across nodes in the tree using the node numbering from the formatted tree for the fixed and variable tree sizes. For the variable size tree, nodes that did not exist in the tree and were zero-padded were not included in the summaries.

To compare the accuracy of inferences with networks trained with fixed or variable tree sizes, ASR was performed on the test dataset from the 50-tip trees using the network trained on trees with 50 tip, between 10 and 50 tips, between 20 and 100 tips, and between 40 and 200 tips. Similarly, ASR was performed on the test dataset from the 100-tip trees using the network trained on trees with 100 tips, between 20 and 100 tips, and between 40 and 200 tips.

### State Speciation and Extinction Models (SSE)

#### BiSSE

Phylogenies were simulated with a BiSSE model (Maddison et al., 2007) using CASTOR. The log 10 birth rates were drawn from unif(−2, 0) for both states. The death rates were drawn from the same distribution and multiplied by the birth rate for the state. This ensured the net diversification rate was positive for each state. The log 10 state transition rates were drawn from unif(−2, 0). The starting state was drawn from the stationary distribution. The maximum taxa was set to 50. If this resulted in full extinction, the parameters were redrawn and the tree was resimulated. As before, ASR was performed with phyddle.

For comparison with Bayesian inference, ancestral states were estimated with RevBayes for the test dataset. The priors were the same as the distributions used in simulation. The prior on the root state was set to the stationary distribution. Two MCMCs were run for each dataset. The MCMCs were run for 15,000 iterations with a burnin of 1,500 iterations and a tuning interval of 100. All moves had weights of 3. To be considered converged, both MCMCs for each dataset were required to have ESS values of 200 for all parameters and the Gelman statistic was required to be at most 1.1 for all parameters. 53 of 2500 datasets were removed due to lack of convergence.

A birth-death model with a Markov model of binary character evolution is special case of the BiSSE model. However, for the Markov model datasets generated in the previous section, the forward and backward transition rates were equal, leading to a more even distribution of tip states than would be generated if the rates were unequal. To compare inferences of ancestral states from a BiSSE model to a Markov model, another dataset was generated using a birth-death model with unequal forward and backward transition rates for a binary character. The simulation set up was similar to the simulations for 50-tip trees. However, both the forward and backward transition rates were drawn independently from the same distribution as was used for the BiSSE simulations, and the root state was drawn from the stationary distribution. Phyddle and RevBayes were used to estimate ancestral states. For RevBayes, the priors matched the distributions used for simulation. The analysis was run for 25,000 iterations. The burnin was an additional 2,500 iterations with a tuning interval of 100. The proposal on rate had weight 2. Convergence was assessed in the same way as it was for the fixed and variable tree size results. 105 of 2,500 datasets were removed due to lack of convergence.

The accuracy of the Bayesian and phyddle estimates were assessed by height by binning nodes according to the proportion of tree height and comparing the proportion of inferences where the inferred state was correct, as described for the fixed tree size for a Markov model of binary character evolution. The probabilities inferred with Bayesian inference and phyddle were also compared.

#### GeoSSE

Phylogenies with 50 tips were simulated with a GeoSSE model (Goldberg et al., 2011) using the R package DIVERSITREE (FitzJohn, 2012). Parameters were drawn from exponential distributions with rate parameter 0.1. All species range states (A, B, AB) were assumed to be equally probable at the root. If full extinction occurred, the parameters were redrawn and the simulation was repeated. The tip state data was recorded as either presence or absence for region A and region B as separate variables. Diversitree always assigns the widespread species to the same daughter node (left vs right) during simulation. This results in the training dataset not containing all possible internal node states, in particular, at the tips where the PHYDDLE formatting step does not rotate branches of equal lengths. To overcome this issue, half of the nodes in the tree were randomly sampled, and those nodes were rotated prior to saving the tree. The encoding of the parent daughter triplet of states was modified accordingly (Fig. 2c). There are eight possible parent daughter triplets with a GeoSSE model with two regions, of which six correspond to rotating the left and right daughters of a different triplet. The ancestral states were estimated as one categorical variable with eight states.

Ancestral states were estimated with RevBayes using TENSORPhylo for 2,500 trees from the test dataset (May and Meyer, 2025). All seven parameters had exponential priors with rate 0.1. All ranges (A, B, or AB) were assumed to be equally likely at the root. The tree was conditioned on time. The MCMCs were run for 50,000 iterations with 5,000 iterations for burnin and a tuning interval of 50. The dispersal and extinction rates moves had weight 1 while the other moves had weight 4. Two MCMCs were run for each data set. All but one pair of MCMCs had ESS values above 200 for all parameters and a Gelman diagnostic below 1.1. One dataset did not meet those criteria, and was removed from the analysis summaries.

### Empirical

#### Liolaemus Lizards

A phylogeny of *Liolaemus* lizard species and their classification of ranges as Andean (highland), non-Andean (lowland), or both was acquired from Esquerre et al. (2019). A subclade of 52 species was extracted for analysis with phyddle based on its size. The ancestral states were estimated with both phyddle and RevBayes using a GeoSSE model. For the GeoSSE analysis in phyddle, the same simulation conditions were used as in the simulations for the GeoSSE model except the tree size was set to 52. A dataset of 500,000 trees was simulated. The batch size was set to 2048.

For RevBayes, the priors for the within-region speciation rates were exponentially distributed with mean 0.1. The same prior was used for the extinction rates, between-region speciation and dispersal. The tree was conditioned on time. All ranges (A, B, AB) were given equal prior probability at the root. The MCMC was run for 15,000 generations with a burnin in 1,500 and a tuning interval of 50 during the burnin. The moves on the speciation rates had weight 4, and the other moves had weight 1. Two replicates were run. Convergence was check by manual inspection with TRACER. All parameters had ESS values above 1000 and the two runs had very similar posterior distributions. The empirical phylogeny was plotted with RevGadgets (Tribble et al., 2022).

#### Ebola virus

Ancestral locations were estimated for sequences from the 2014 West African Ebola virus outbreak (Dudas et al., 2017) using only a subset from Sierra Leone (Müller et al., 2019). The MCC tree and location data from Müller et al. (2019) were downloaded. The tree was randomly subsampled to 50 sequences, negative branch lengths were changed to be small positive values, and branch lengths were converted into units of weeks. This included sequences sampled during May to December of 2014, which was past the time of peak incidence for most of the 14 districts in Sierra Leone. Based on incidence data, the outbreak in Sierra Leone was likely to have started in Kailahun. We assigned each viral sample to 1 of 5 regions, where our regions were agglomerations of adjacent regions from the original 14-region system of Müller et al. (2019) (see their Fig. 3b). Our location 0 included Kailahun, Kenema, and Bo; location 1 included Kono, Koinadugu, and Bombali; location 2 included Tonkolili, Port Loko and Kambia; location 3 included Pujehun, Moyamba, and Bonthe; and location 4 included the Western Rural and Western Urban regions. In addition to the phylogenetic tree and sequence locations, we also provided the regional times of peak prevalence as input to help phyddle constrain when the disease probably entered a population. The time of the first case in Sierra Leone in the incidence data was treated as week 1. The times of peak prevalence for each region were determined from the incidence data by assuming a two week infectious period and summing the incidence data for the previous two weeks (Qureshi et al., 2015; Legrand et al., 2007). The times of peak prevalence were very similar if an infectious period of one week was assumed.

Ancestral states were inferred with phyddle using a SIR + Migration (SIRM) model with five regions. A dataset of 500,000 phylogenies was simulated (Riley, 2007) using MASTER (Vaughan and Drummond, 2013) with custom python scripts to produce the xml files. The number of hosts in each location was set to 10^4^. The basic reproductive number, *R*_0_, was drawn from unif(1, 4). The recovery rate was drawn from unif(10^−2^, 4 ×10^−2^). The sampling rate was drawn from unif(10^− 1^, 1). The migration rates were drawn from unif(10^−5^, 4.9 ×10^−3^). The stopping time was set to 50 weeks. The maximum number of sampled taxa was set to 50. The probability that an extant tip was sampled was 10^−4^. The incubation period was not explicitly modeled. Regional times of peak prevalence were generated by MASTER as part of simulation, and were provided to phyddle for training. The neural network was trained fives times, where we used the mean state probabilities across the five networks to produce our main results. Results based on each individually trained network are shown in the supplement. The empirical phylogeny was plotted with RevGadgets (Tribble et al., 2022) and GGTREE (Yu et al., 2017).

### Phyddle *configuration*

Phyddle uses convolutional and pooling layers and dense feed-forward layers to construct its neural network (Landis and Thompson, 2025). All analyses used the same settings, unless stated otherwise. The default settings for phyddle were used for the network architecture (e.g. numbers of layers, numbers of node per layer, etc.). For each analysis, 250,000 trees were generated for each experiment, with 95% used for training and 5% used for testing. Of these, only 2,500 of the trees from the test datasets were used for generating the figures and results, and test dataset is used to refer to only those 2,500 trees. This allowed us to match the trees used in the Bayesian analyses, which also used 2,500 trees. The tree width was set to the maximum size of the trees in the training dataset except for the four-tip trees. For the four-tip trees, the tree width was set to 32, since lower tree width settings would have required the default values for kernel to be adjusted. Characters were encoded with integer encoding and the trees were encoded as extant only trees, with the exception of the SIRM model which used serially sampled trees. The brlen encode setting was assigned to height brlen, which includes additional branch length and node age information in the tree tensors in comparison to the original CDV or CBLV format. This additional information is shown in Fig. S1c.

## Results

### Markov model of binary character evolution

#### Four-tip trees

ASR was performed on four-tip trees using Bayesian inference and three estimation strategies with phyddle. The marginal strategy estimated the marginal probabilities of each of the three nodes using a single network that estimated three variables. The joint strategy estimated the probability of each possible combination of the three internal node states jointly with a single variable. The single node strategy estimated the marginal probability of one node using a network that takes as input the node name and only estimated the marginal probability of the state at that node. These strategies were compared against Bayesian estimation of ancestral states.

The most probable state was considered the point estimate for the marginal and single node estimation strategies as well as for Bayesian inference. For the joint estimation strategy, the probabilities were marginalized to compare to the other methods. The proportion of inferences where the most probable state was correct was very close across all methods (Table 1 diagonal). The frequency with which the inferred state from one method matched the inferred state from a different method was high across all method pairs (Table 1 off diagonal). The probabilities of the ancestral states were similar for phyddle and Bayesian inference (Fig. S2).

**Table 1.** Proportion of inferences where the point estimates (most probable states agree) of ancestral states that agree between methods for the test dataset (off diagonal). The diagonal is compared with the true ancestral state. The first three columns and rows are with phyddle and the last is with Bayesian inference. The matrix is symmetric, so only half is filled in.

|  | joint | marginal | single node | Bayesian |
| --- | --- | --- | --- | --- |
| joint | 0.82 | 0.93 | 0.92 | 0.94 |
| marginal |  | 0.82 | 0.93 | 0.93 |
| single node |  |  | 0.82 | 0.95 |
| Bayesian |  |  |  | 0.83 |

For a symmetric four-tip tree, there are eight possible combinations of internal states (Fig. 3a), and similarly there are eight combinations for a four-tip asymmetric tree (Fig. S3a). Some of these combinations may be easier to infer, such as when the minimum possible number of changes on the tree is smaller. The proportion of inferences where all three internal nodes were estimated correctly for a given combination of true internal node states was similar across the three estimation strategies with phyddle (Fig. 3b). This proportion varied between trainings, sometimes over 0.10 for a given method and combination of ancestral states. For the pairs of trees with the 0 and 1 assignments switched (e.g the columns in Fig. 3a), the proportion of correct inferences was similar, but not the same. It may be expected that using zero-padding could cause a bias toward state 0, so trees with more 0s would be correct more often. However, there was not a consistent pattern in which trees had a higher proportion of correct inferences. As expected, as the minimum number of changes required in the true history increased, the accuracy of the inferences decreased. The patterns were similar for four-tip trees with asymmetric topologies (Fig. S3).

#### Fixed tree size

The accuracy of the probabilities of the ancestral states estimates was assessed for each of the three fixed tree sizes. Comparing the proportion of correctly inferred states to the probabilities of the state from phyddle, the ancestral state probabilities were fairly accurate on average (Fig. S4). The probabilities were more accurate at when they were close to 0, 0.5, or 1. They were slightly too high if they were between 0 and 0.5 and slightly too low if they were between 0.5 and 1. The 200-tip trees had the most accurate probabilities, with the probabilities reported by phyddle being closest to the proportion of inferences where the most probable state was correct on average. The proportion of the root nodes where the most probable state was the correct state was 0.70, 0.67, and 0.66 for the 50, 100, and 200 tip trees using the marginal estimation strategy. The proportion correct was slightly higher when estimating the root node age as a single variable using the approach of Thompson et al. (2024), at 0.71, 0.71, and 0.69 for the 50-, 100-, and 200-tip trees, respectively.

#### Comparison of fixed and variable tree size

The proportions of inferences where the point estimate was correct, when averaged across all nodes in the test datasets for each fixed size tree, were 0.80, 0.78, and 0.72 for the 50-, 100-, and 200-tip trees, respectively. The proportions were comparable for the variable tree sizes (0.80, 0.78, and 0.74 for the trees of up to 50, 100, and 200 tips, respectively). The ancestral states for the shallow nodes were more accurately inferred than deeper nodes in the trees (Fig. 4a). Accuracy was lowest for 200-tip trees. The accuracy of Bayesian inference was higher than phyddle across all tree sizes. The discrepancy between the accuracies of the two methods grew as the tree size increased. The results were very similar with variable size trees (Fig. S5).

**Fig. 4.**
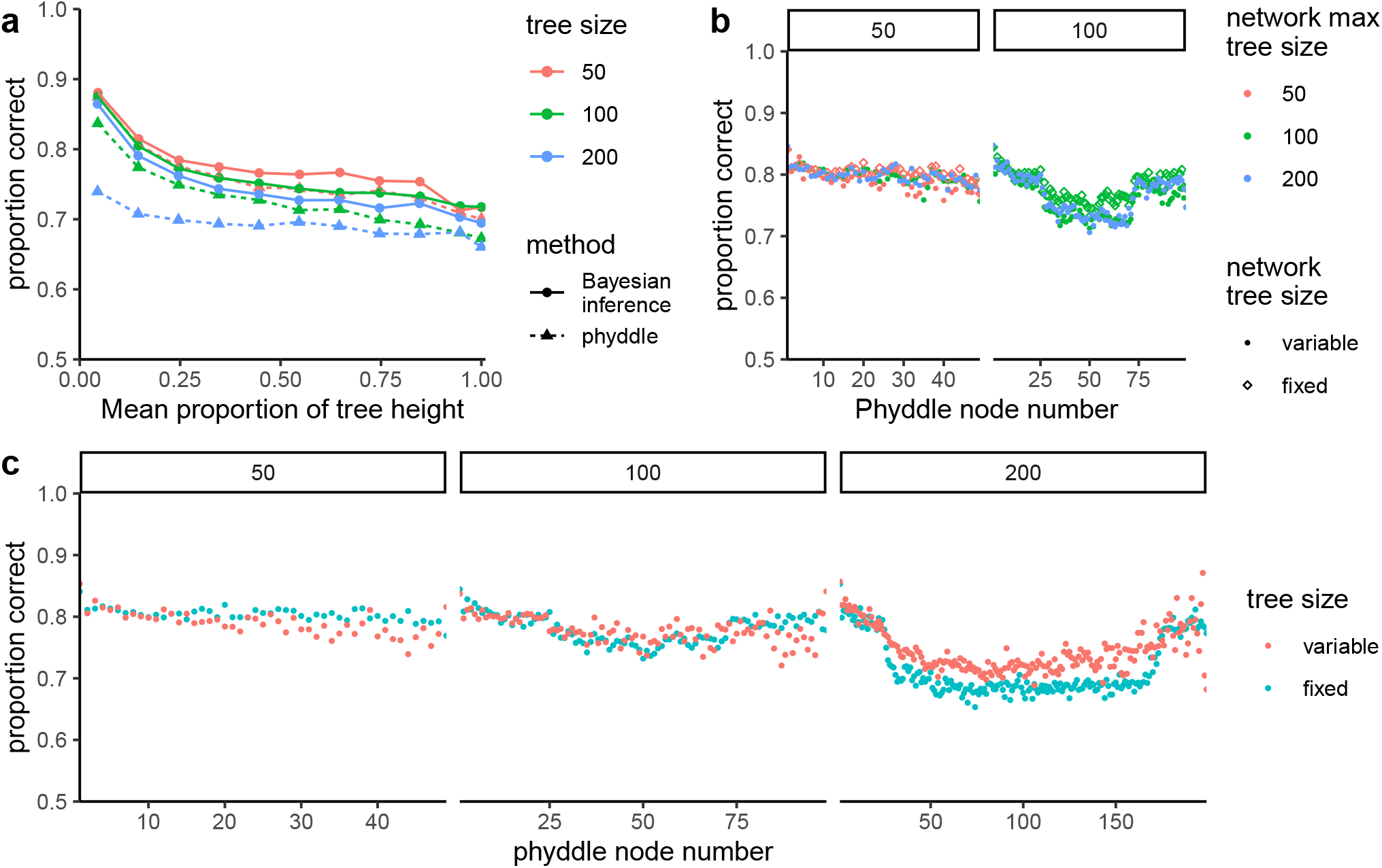
(a) The proportion of inferences where the inferred state was correct by node height relative to tree height for fixed tree sizes. The colors show the tree size and the line and dot types show the method used for inference. (b) The accuracy of inferences for trees of a fixed size on networks trained with fixed or variable tree sizes. The colors show the maximum tree size seen by the network in the training data. The shapes show whether the network was trained with fixed or variable size trees. The panels show the performance based on tree size. (c) The average accuracy by node for fixed and variable tree sizes. The panels show different maximum tree sizes and the colors show inferences with fixed and variable tree sizes using test datasets simulated in the same way as the training datasets. The node number corresponds to the node number in the tree formatted by phyddle. The y-axes of all panels range from 0.5 to 1.0.

Different nodes in the formatted tree correspond to different average positions within the tree. For example, the lowest node number always had two tips as daughters. With variable size trees, the highest node number had the fewest examples in the training dataset. Comparing the accuracy of individual nodes, the smallest node numbers tended to have the highest accuracy for a given tree size (Fig. 4c). Larger node number tended to have lower accuracy in the variable size trees in comparison with the fixed size trees. The 200-tip trees had lower accuracy overall, with the middle node numbers having higher accuracy for the variable trees in comparison to the fixed size trees. This could be due to those nodes corresponding to smaller trees on average for the variable tree size inference, making the inference problem easier.

When the performance of neural networks trained on different tree sizes were evaluated with a single tree size, the performance did not vary much across the range of tree sizes in the training data (Fig. 4b). For inferences on 50-tip trees, the proportions of point estimates that were correct were 0.79, 0.80, and 0.80 for networks trained on trees up to 50, 100, and 200 tips, respectively. For inferences on 100-tip trees, the proportions correct were 0.76 and 0.77 for networks trained on trees up to 100 and 200 tips, respectively. The proportions correct were slightly higher when using a network trained on a single tree size compared to variable size trees, 0.80 and 0.78 on average for 50 and 100 tip trees, respectively.

### State Speciation and Extinction Models

#### BiSSE

Inferences of ancestral states using phyddle for BiSSE and Markov models had similar proportions of point estimates that were correct for shallow nodes in the trees, with ancestral state states inferred correctly slightly more often with a BiSSE model (Fig. 5a). For deeper nodes in the tree, the accuracy of both Bayesian inference and phyddle declined and was lower for the Markov model. The difference in the probability of state 1 estimated by phyddle and Bayesian inference was typically quite small for both the BiSSE and Markov models (Fig. 5b). However, on rare occasions the difference between the probabilities was quite large (tails in Fig. 5b). The probabilities reported for state 1 by phyddle and Bayesian inference were strongly correlated, for both models (Fig. 5c). The correlations for the Markov and BiSSE models was 0.98 and 0.95, respectively. 95% of all inferences fell between the between the dashed lines (Fig. 5c). However, there was much more noise than in the simpler case of a Markov model with symmetric rates on four-tip trees (Fig. S2). phyddle outperformed a naive estimator which estimated the ancestral state at each node to be the most common state of its descendants (Fig. S6).

**Fig. 5.**
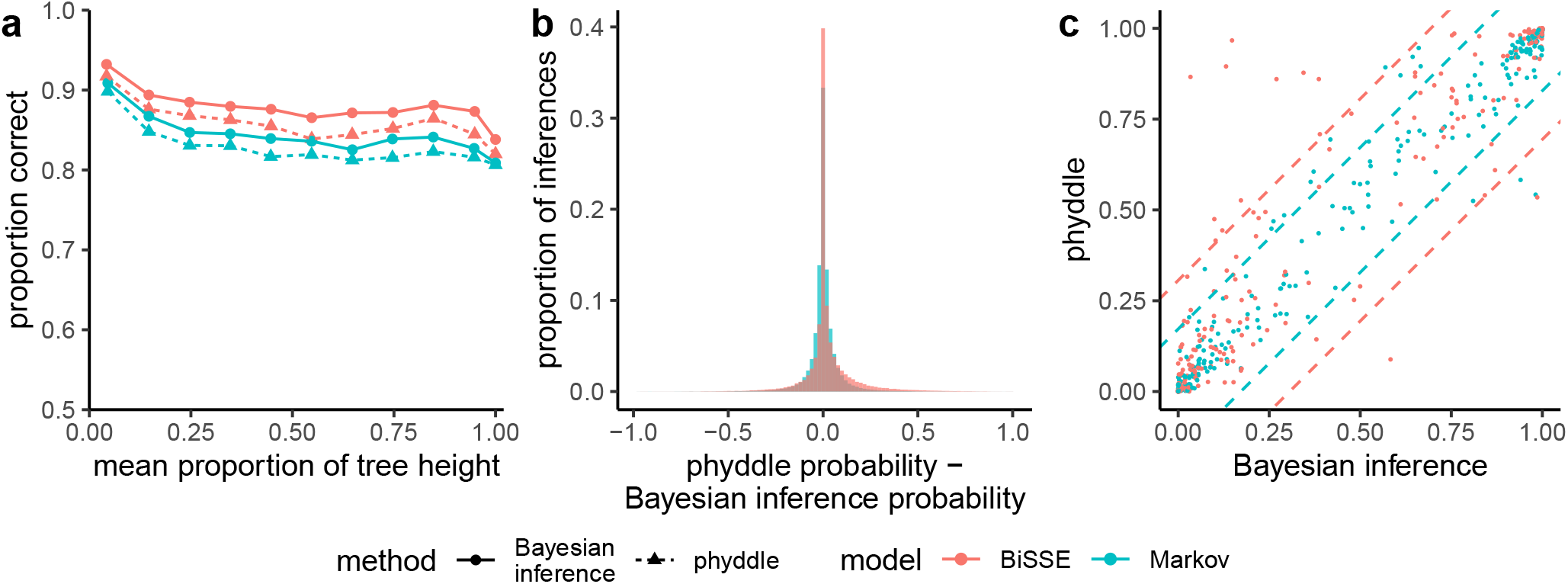
(a) The proportion of inferences where the inferred state was correct by the proportion of the tree height comparing Bayesian inference and phyddle for a BiSSE and Markov model of character evolution. The y-axis ranges from 0.5 to 1.0. (b) The difference between the probabilities of ancestral state 1 estimated by phyddle and Bayesian inference for all nodes in the test dataset. (c) The probability of state 1 from phyddle and Bayesian inference for one node for 500 randomly sampled trees in the test dataset. 95% of the inferences fall between the dashed lines of the corresponding color for each model.

#### GeoSSE

For a GeoSSE model, the parent-daughter triplets of ancestral states were inferred as a categorical variable with eight possible states. When the true ancestral range was a single region (A→A,A or B→B,B), the proportion of point estimates inferred by phyddle that were correct was high (Fig. 6). The proportion was lower if the parent node was widespread (AB). However, the errors were not random and were similar to the errors from Bayesian inference. For example, a widespread parent that gave rise to one widespread daughter and one daughter that occupied only region A (AB→AB,A or AB→A,AB) was often inferred to have the parent and daughters all occupying region A alone. A similar pattern arose for when one of the daughters was only in B (AB→AB,B or AB→ B,AB). The accuracy was similar for the states where the daughters were rotated (e.g. AB→A,B and AB→B,A). The overall proportions of inferences where the most probable state was correct were 0.640 and 0.652 for phyddle and Bayesian inference, respectively. Phyddle had higher accuracy than Bayesian inference when the true parent state was a single region, but lower accuracy than Bayesian inferences for all cases when the parent was widespread. When one of the daughters was widespread, the accuracy of inference with phyddle was lower than when the parent was widespread but neither daughter was widespread. On average the probabilities of the states A→A,A and B→B,B were slightly higher when estimated by phyddle than when estimated with Bayesian inference (Fig. S7). When there was a between-region speciation event, the mean difference in probabilities was very small. For states with widespread daughters, the Bayesian probabilities were slightly larger than the phyddle probabilities on average.

**Fig. 6.**
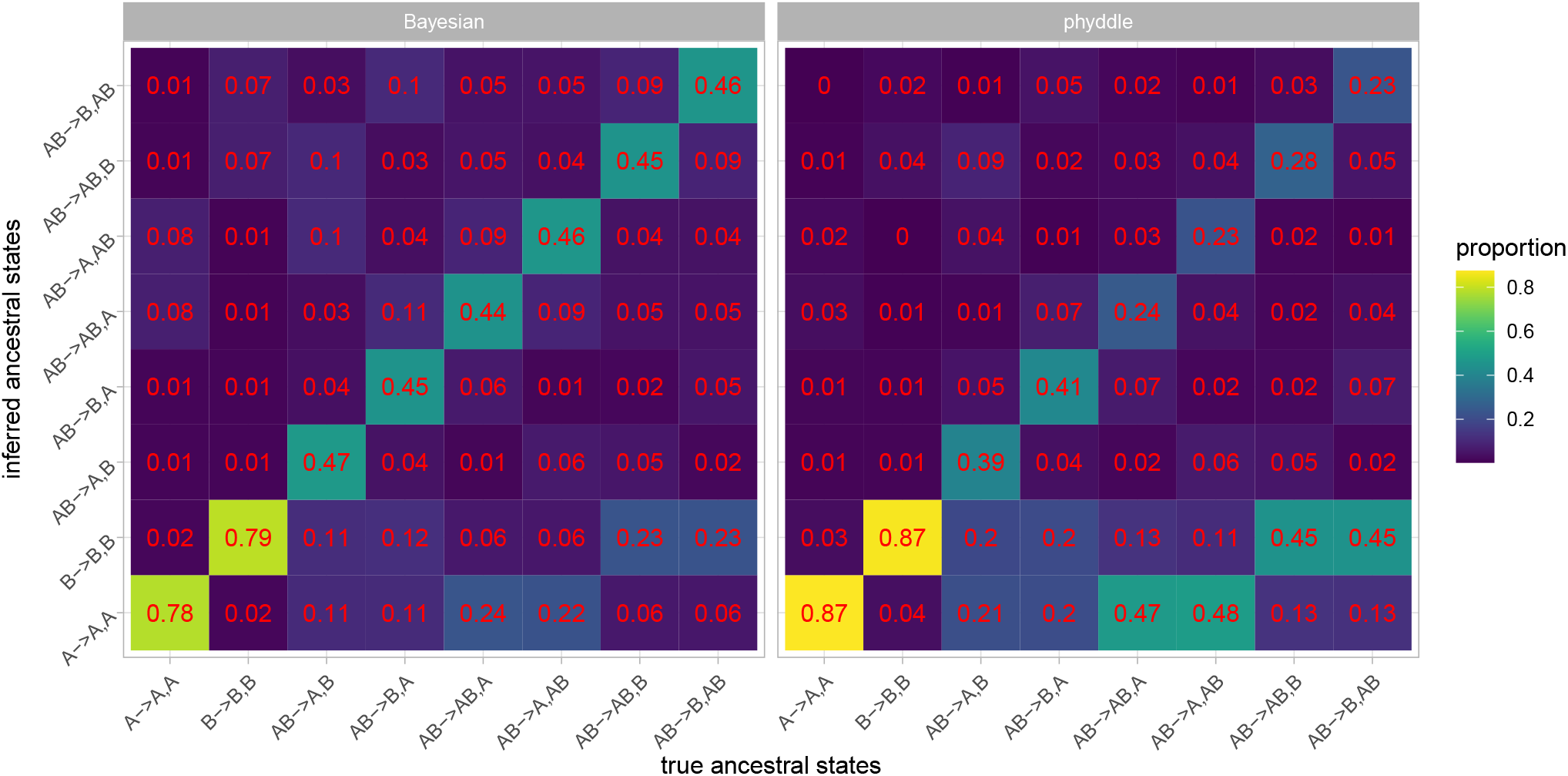
Confusion matrix for GeoSSE inferences using Bayesian inference (left) and phyddle (right) as a heat map. The columns show the true ancestral state. Within a column, the rows show the proportion of inferences where the ancestral state on the y-axis is preferred (i.e. has the highest probability). The proportions within a column sum to 1. The numbers on each square match the coloring of the heat map.

### Empirical

#### Liolaemus Lizards

The ancestral state reconstructions under a GeoSSE model as inferred by phyddle and Bayesian inference were similar (Fig. 7 and S8). The inferences tended to be highly concordant in regions of the tree with little variation in tip states among the daughter nodes, with most discordance deeper in the tree. Phyddle inferred higher probability of Andean ancestral ranges deeper in the tree than with Bayesian inference.

**Fig. 7.**
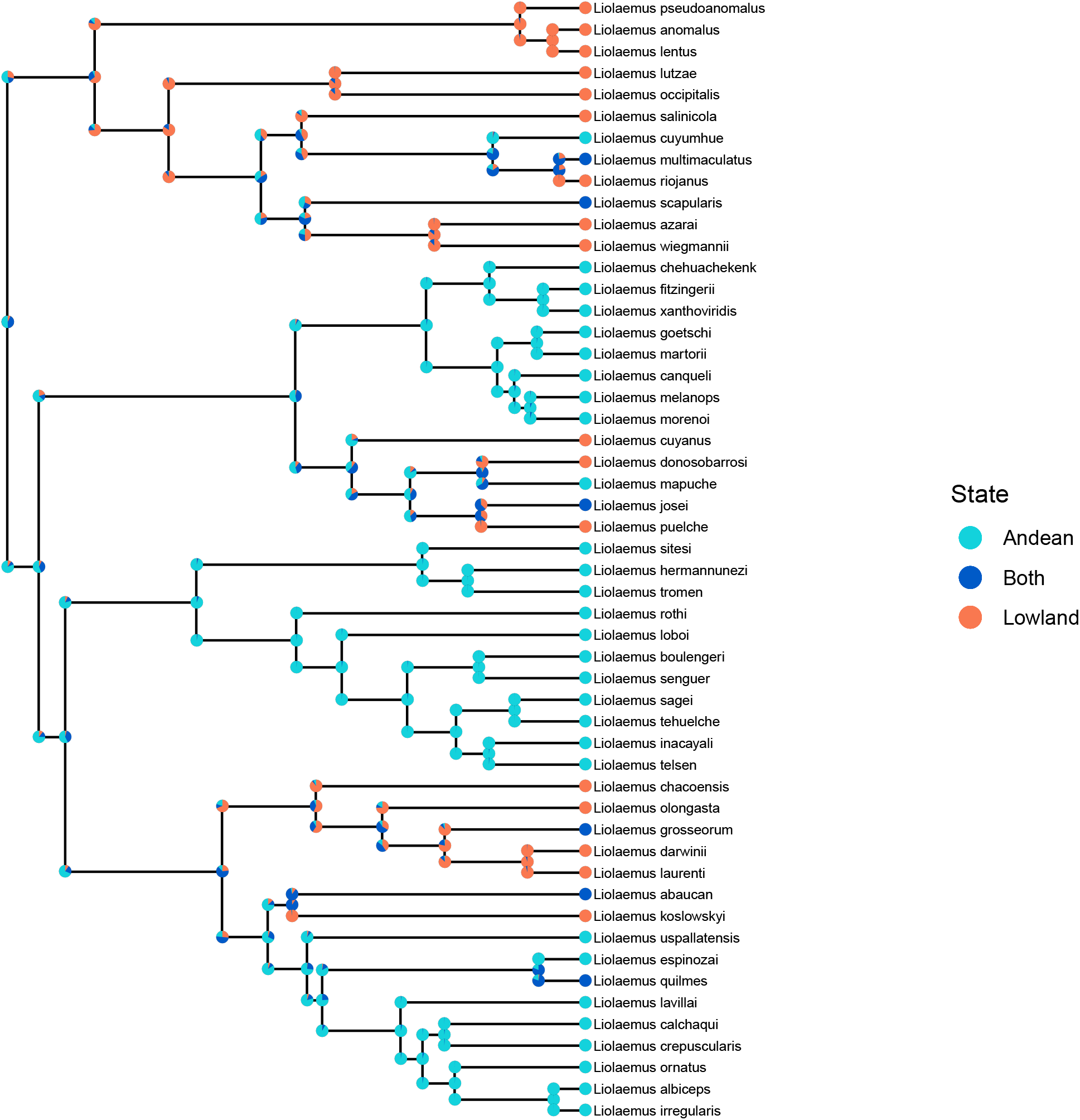
Ancestral state reconstruction for a subclade of 52 *Liolaemus* species with a GeoSSE model using phyddle. The pie charts centered at the internal nodes show the probability of the parent state at the time of speciation. The pie charts on the daughter branches show the probabilities of each of the daughter states immediately after speciation. These probabilities were found by marginalizing over the 8 possible parent daughter triplets of possible states.

#### Ebola virus

The accuracy of the ancestral state estimates in the SIRM test dataset was high; the proportion of inferences where the most probable state was the true state was 0.95. Some of the summary statistics for the empirical Ebola tree were out of distribution in comparison to simulations, including colless, N-bar, and treeness as calculated by Dendropy within phyddle (Moreno et al., 2024). The most probable ancestral state reconstructions for the locations of the Ebola under a SIRM model as inferred by phyddle were largely consistent with what one would expect from a parsimony-based reconstruction. Nodes with little variation in the daughter states tended to have the favored ancestral state be inferred with high probability (Fig. 8). The deepest nodes in the tree were inferred to be in state 0 and were inferred with high probability, consistent with the epidemiological data. However, a few nodes in the tree were inferred to be in state 2, in some cases despite having no descendant nodes in state 2. In these cases, the second most probable state matches one of the descendants (Fig. 8). Different trainings of the neural network gave slightly varying results (Fig. S9-13), but uncertain nodes tend to remain uncertain and strongly supported nodes tended to remain strongly supported.

**Fig. 8.**
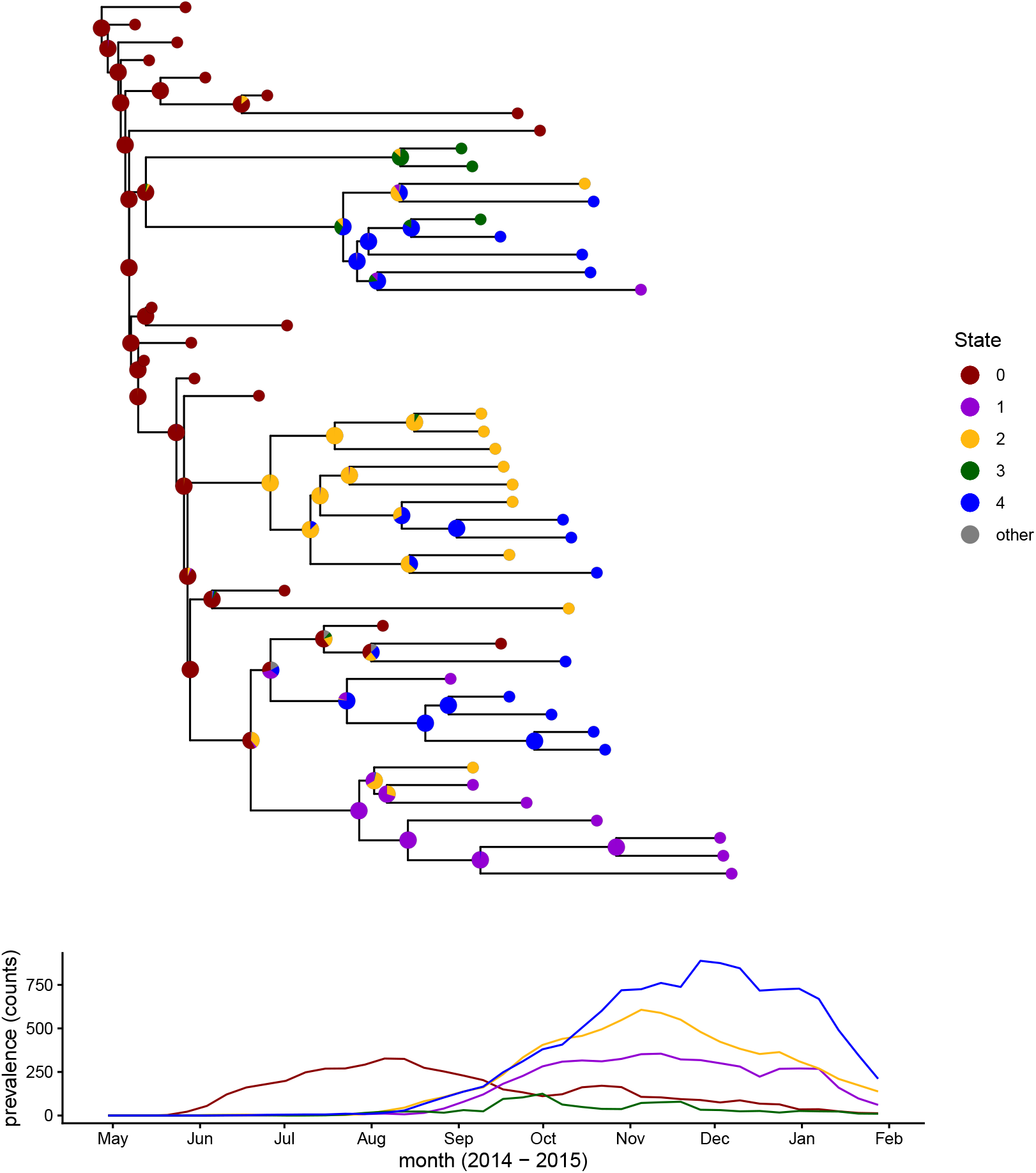
Ancestral location reconstruction for Ebola viruses sequences from the 2014 outbreak in Sierra Leone with the prevalence by region. The pie charts show the probabilities of each state (region) inferred by phyddle. Locations correspond to groups of districts within Sierra Leone. State 0 corresponds to Kailahun, Kenema, and Bo. State 1 corresponds to Kono, Koinadugu, and Bombali. State 2 corresponds to Tonkolili, Port Loko and Kambia. State 3 corresponds to Pujehun, Moyamba, and Bonthe. State 4 corresponds to Western Rural and Western Urban. Only the three most likely ancestral states are shown by the state color. Any remaining probability for the other two states is shown in gray. The prevalence over time is shown for each population. The prevalence was calculated by summing the incidence from the previous two weeks with the current week. The time axis is aligned for the plots.

## Discussion

The goal of our study was to introduce a simple method for ASR using deep learning and rigorously evaluate its performance so it can be used as a benchmark for future studies. We explored three alternative strategies for estimating ancestral states (marginal, joint, and single node), and explored how to estimate triplets of ancestral states for models that support asymmetric cladogenetic state inheritance patterns. Our method’s accuracy resembled that of Bayesian inference in several cases, particularly for small trees (~50 or fewer taxa). Accuracy tended to decline as the tree size increased, when the training datasets were too small, and/or when model complexity increased. To assess the method’s performance under empirical settings, where the true data-generating model is not known, it was applied to the historical biogeography of *Liolaemus* lizards (Esquerre et al., 2019) and to the 2014 outbreak of Ebola in Sierra Leone (Müller et al., 2019). For the latter, an SIR model with host migration was used, for which it is not trivial to compute the exact likelihood of generating a particular time-calibrated tree of viral sequences, sequence locations, and times of peak prevalence per region under a set of unknown model parameters.

For small trees with a Markov model, the accuracy of phyddle and Bayesian inference were similar, independent of which deep learning strategy was used. All methods tended to infer the same ancestral state, and there was a strong correlation between the probabilities inferred by phyddle using any of the three strategies and Bayesian inference. Accuracy for specific trees varied substantially for different trainings despite overall accuracy remaining similar. As the tree size increased using a Markov model, phyddle’s performance relative to Bayesian inference dropped, and deeper nodes in the tree were inferred less accurately. Networks trained on different distributions of tree sizes performed similarly to each other given the same test dataset, suggesting that generalizing across tree sizes was not a major obstacle to learning. The increase in topological complexity may have been a larger factor in the decrease in performance for larger trees. There were no obvious or consistent biases in preference for 0 or 1 for any of the experiments with Markov models, which could occur if zero-padding for variable size trees biases the inference.

For SSE models, similar patterns held. For a BiSSE model, phyddle was more accurate for shallow nodes, less accurate for deeper nodes, and the probabilities from deep learning and Bayesian inference were strongly correlated. However, occasionally the probabilities reported by phyddle substantially differed Bayesian inference, and phyddle was confidently wrong. In our analyses, while the overall proportion of nodes where the most probable state was correct was very similar for phyddle and Bayesian inference for a GeoSSE model, the errors were distributed differently for the two methods. phyddle had a preference toward inferring single region states in comparison to Bayesian inference. Single region ancestral ranges were the most common in the training data, which may have contributed to estimates that were biased for single-region reconstructions. One possible solution is to reweight the cross-entropy loss score to place greater importance on correctly inferring rare states.

For the *Liolaemus* analyses, phyddle’s inferences appeared reasonable, and often agreed with the Bayesian estimates, but with some exceptions. For example, the node that is ancestral to *Liolaemus abaucan* and *Liolaemus koslowskyi* was inferred to have a range of both Andean and lowlands. Its parent node is separated by only a short branch length and inferred to have a single region as the range. This requires gaining a region on a very short branch, rather than gaining the region on the much longer branch to *L. abaucan*. Disagreements between phyddle and Bayesian inferences most often manifested among deeper nodes and for nodes with more variation among the tip states of their descendants. Despite such oddities in this empirical example, the performance of phyddle and Bayesian inference were quite similar for the simulated test data. Assuming the simulation conditions were sufficiently general, it is reasonable to believe that both sets of inferences are equally accurate in the point estimates, on average.

In the location reconstructions for the Ebola virus sequences, the deeper nodes in the tree were confidently inferred to have occurred in region 0, reflecting what the infection prevalence data portray. Many shallower nodes, particularly when all sampled descendants shared a location, also produced confident reconstructions that were plausible. However, the estimates for some nodes did not intuitively make sense; locations that were not represented in the daughters were inferred as the ancestral location for a couple nodes. Additionally, some ancestral states were inferred prior to a rise in prevalence in the corresponding regions. This could be partially due to a delay in or under reporting of cases, especially early on in the epidemic (Dalziel et al., 2018) or due to undetected transmission of asymptomatic or mildly symptomatic cases (Richardson et al., 2016). Inferences with complex models, such as an SIRM model, required larger training datasets to perform well, as smaller datasets yielded worse loss scores (results not shown). This may limit the types of models that a researcher can use, depending on their access to computational resources. We also found that different, independent trainings of the network yielded somewhat different results, motivating us to use the average predictions across all networks. These results suggest that more work is needed to better capture the patterns in complex models for ancestral state reconstruction.

A general challenge for training neural networks in phylogenetics is simulating realistic training datasets. In the context of ASR, naively choosing parameter combinations can unpredictably lead to very little variation in the tip states or very high rates of change between parent and daughter nodes, neither of which is guaranteed to represent an arbitrary empirical system that a biologist may want to analyze. With more complex models, such as SSE or SIR models, it can be difficult to choose parameters that generate the desired amount of variation, in terms of traits, taxa, and timescales. Training datasets that are not representative of empirical data may lead to biased inferences and lower performance on empirical data than simulated data. For example with a GeoSSE model, one region may have a very small, or even negative, net diversification rate, as may have been the case for the chaparral plant clades analyzed by Goldberg et al. (2011). However, to produce trees with extant taxa under such conditions, it also requires that the species originate in a second region that has a positive net diversification rate and a high dispersal rate. Targeting this relatively constrained part of parameter space (and pattern space) during simulation is not always trivial, and these issues are exacerbated for more complex models where likelihood-based inference is not possible. We did not obtain a realistic simulated training set for our empirical analyses on the first try. Rather, we combined mathematical reasoning, biological knowledge, exploratory analyses, and out-of-distribution tests (provided with phyddle) to experiment with our simulation design before finding conditions that better resembled our empirical systems.

Moreover, training datasets can be biased even if the underlying simulation methods are correct and cover a large range of parameter space. On one hand, this can be because the simulator is misused, and fails to represent the intended simulation scenario, e.g. accidentally assigning the root node to the same state across all simulations. On the other hand, the simulator might make strong and non-random assumptions when recording data patterns to computer memory. For instance, when there is a within-region speciation event under the GeoSSE model, DIVERSITREE always assigns widespread species and single region species to the same daughter branches (left versus right). Automatically designating every widespread daughter lineage as (e.g.) the left node generally will not impact a well-designed likelihood function. However, for deep learning this leads to a lack of all possible patterns in the training data, where there is no reason to expect this pattern in empirical data, as it only depends on the rotation of the internal nodes in the tree. Other simulators may have similar default behaviors that lead to poor outcomes in the inferences for empirical data. Unfortunately, in some cases preventing these issues may require knowledge of the implementation details of software, or at a minimum checking for unexpected patterns in the simulated data. As such, users of deep learning should be cautious when using existing simulation software to train neutral networks.

A major difficulty in ASR with deep learning is generating training data with sufficiently representative patterns so that the neural network can infer ancestral states for nodes in topologies the network has never seen. Moreover, even if there are nodes that are partially comparable across topologies, such as nodes within subclades with matching topologies, the data need to be provided in a way that the neural network can learn patterns. phyddle uses a relatively standard architecture, comprised of convolutional and pooling layers for processing the tree tensors, and dense layers for processing the auxiliary data, which we also used for our study. That said, this architecture was initially designed for generic inference problems using phylogenies, such as parameter estimation or model selection (Landis and Thompson, 2025). Such tasks should be easier for neural networks as the same parameters exist for every tree, as opposed to ASR since same nodes do not exist for every tree and the number of states to infer scales with the tree size. Given the simplicity of ASR method, it is surprising that phyddle achieves reasonable results in many cases.

Neural network architecture is known to be important in deep learning performance. The number of layers, kernel settings (width, stride, dilation), batch size, and encoding type may all alter method performance. The default settings for phyddle appeared to work reasonably well in our experiments. A deeper architecture with additional convolutional layers may improve performance, particularly for larger trees. Users should consider experimenting with the settings, particularly if they experience poor performance or do not compare against a likelihood-based method that can serve as a baseline. Different neural network architectures, such as graph neural networks (Qin et al., 2026; Leroy et al., 2025) may enable better learning of the patterns by explicitly using the tree structure. In particular, explicit use of the phylogeny’s graphical structure should increase the level of correlation among inferred states for adjacent nodes in the tree, and therefore reduce the chance that multiple (e.g. non-parsimonious) changes are required to explain state transition patterns, as occurred in the Ebola dataset.

Formatting the phylogenetic data differently (Perez and Gascuel, 2025) or including additional summary statistics (Janzen and Etienne, 2024), such as tree-based summary statistics targeted toward ASR, may also improve performance. Another strategy may be to first estimate the rate parameters with deep learning, and then provide those estimates as input to another neural network that estimates ancestral states. That said, based on a few number of exploratory experiments, this did not improve our results, but it may help in other instances. For SIR models, outside information such as incidence or prevalence may be informative about ancestral states and could be used as additional input data. In our case, we found that simply adding the regional times of peak prevalence improved ancestral state estimates.

Whenever possible, we compared phyddle’s performance to that of Bayesian inference and generally assumed that the Bayesian inference gave the “right” answer. Even in cases where there is a likelihood-based method with which to compare, the most appropriate comparisons are not always obvious. Any method can give inconsistent results for both statistical and numerical reasons. Maximum likelihood and Bayesian inference may produce quite different results when the data are not informative, e.g. due to non-identifiability and/or prior sensitivity. Methods that implement a maximum likelihood search under the same model do not always give consistent results (e.g. Lambert et al., 2023), and Bayesian methods can suffer from issues such as lack of convergence. Both methods can struggle with numerical integration in some regions of parameter space. In our experiments, the simulation conditions always matched the priors for Bayesian inference and most of the priors were bounded, which should be favorable toward Bayesian inference. More generally, a thorough and fair assessment of deep learning methods must be considered carefully in relation to the assumptions of alternative methods (e.g. maximum likelihood or Bayesian inference).

## Conclusion

It is uncontroversial to say that all phylogenetic estimates of biological data are imperfect, where biologists generally seek to minimize the error in modeling the “truth”. Both the model and the method will each contribute a distinct part to the total estimation error. Likelihood-based methods are expected to have lower method error than deep learning-based approaches when the model is correct. However, many models that increase the biological realism in comparison to standard models for inference do not have tractable likelihood functions. In these cases, the models used under a likelihood-based method decrease method error at the expense of model error. Deep learning methods may, under certain conditions, have a higher baseline level of method error, but their intrinsic compatibility with biologically realistic models help minimize model error. Thus, to evaluate method performance, it is necessary to not only use the standard models for which likelihoods have been derived, but also to consider more biologically realistic models that lack likelihood functions. Future work measuring the relative contribution of estimation error across a variety of models and methods would help the phylogenetics community prioritize research efforts. Even though our deep learning method for ancestral state estimation is far from an ideal solution to the problem, we hope it stimulates further discussion about ways to improve phylogenetic model and method design.

## Acknowledgments

We would like to thank the Landis lab members (Albert Soewongsono, Ammon Thompson, Sarah Swiston, Sean McHugh, Raymond Castillo, and visiting scholar Gabriel Santos Garcia) and Ixchel González-Ramírez for helpful feedback concerning this project. We would also like to thank Nicola Müller for assistance with the Ebola dataset.

## Funding

This work was funded by the National Science Foundation (NSF Award DEB-2040347) and the Fogarty International Center at the National Institutes of Health (Award Number R01 TW012704) as part of the joint NIH-NSF-NIFA Ecology and Evolution of Infectious Disease program.

## Software and Code Availability

Analysis scripts are available at https://github.com/nage0178/asr_phyddle.git. The version of phyddle used in this manuscript is available at https://github.com/mlandis/phyddle/tree/devAncState. RevBayes version 1.4.0-preview commit 10c322 was used for all analyses except the GeoSSE analyses. The GeoSSE analyses used commit 6a276d with TensorPhylo commit 8887a4 (May and Meyer, 2025).

## Notes

### Competing Interest Statement

The authors have declared no competing interest.

