## Supplemental material for "Ancestral state reconstruction with discrete characters using deep learning"

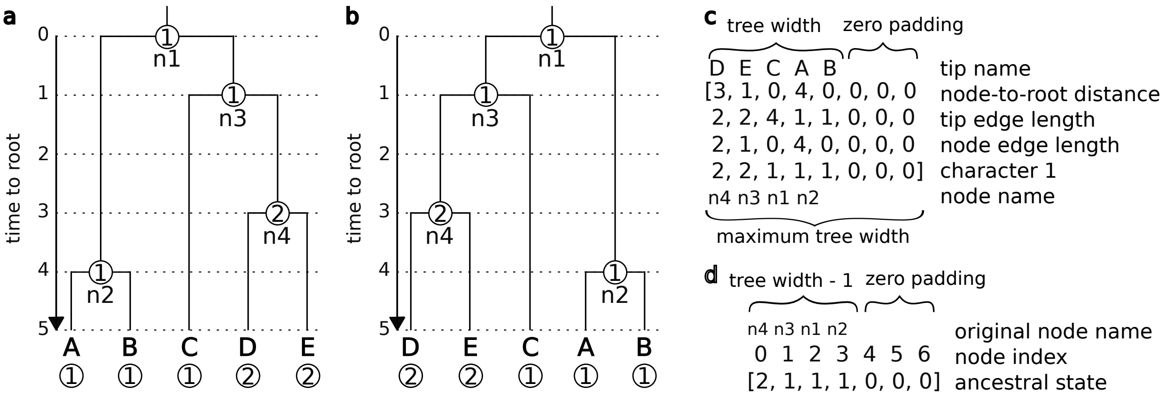


Figure S1: Formatting of the phylogeny tensor and the ancestral states. (a) The input phylogenetic tree has 5 tips and 4 labeled internal nodes (n1, n2, etc.). Each node has node has an ancestral state (1 or 2), which is given in a csv file for training. The tips also have states (1 or 2). (b) The internal nodes in the tree are rotated such that the sum of the branch lengths on the left subtree is greater than the sum of the branch lengths of the right subtree. (c) The tree tensor includes the node-to-root distances, the tip edge lengths, the node edge lengths, and the tip character data. Additional character data at the tips results in additional rows in the tensors. The order of the tips and internal nodes is determined by an in-order tree traversal. Zeros are added to the end of each row when the tree size is smaller than the maximum tree size. Note that PHYDDLE represents each tree as a column, as oppose to rows as is shown for clarity. (d) The ancestral states are provided in a separate tensor as training data. Zeros are added to the end of the tensor when the tree size is smaller than the maximum tree width. Note that for a categorical variable with $n$ states, PHYDDLE changes the labels to be from 0 to $n-1$. This is not shown in the figure for clarity.

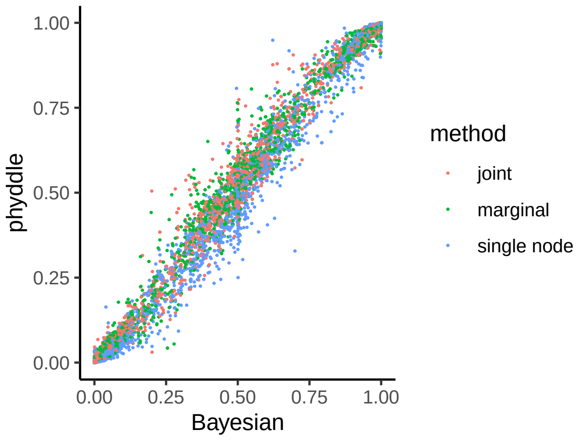


Figure S2: The probability of state 1 as inferred by PHYDDLE and Bayesian inference on four-tip trees. One random internal node is shown from each of 2,500 trees in the test dataset for each method. All three methods use the same set of trees.

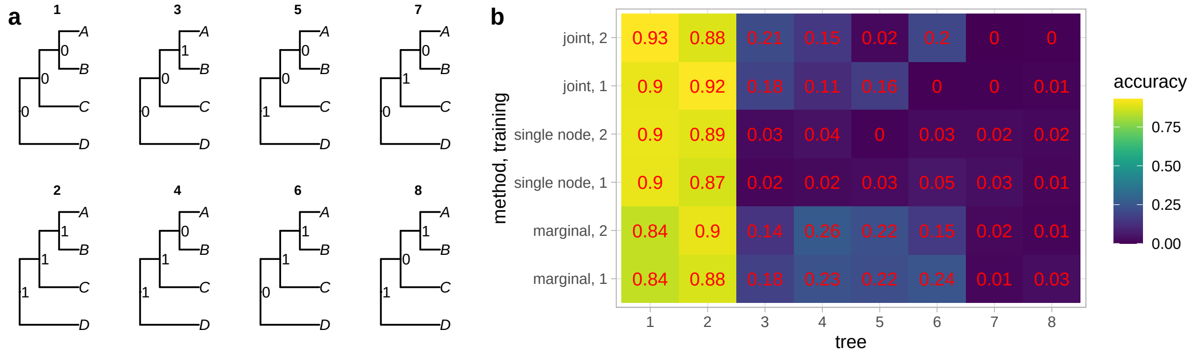


Figure S3: (a) There are 8 possible internal node state patterns for an asymmetric four-tip tree with a binary character, shown in trees 1-8. In the first column, no changes are required on the internal branches to observe the ancestral state pattern. In the second column, at least 1 change is required to observe the pattern of ancestral states and the change is on the branch between C and the AB clade. The third column also requires at least one change on the branch connecting D to the rest of the tree. The last column requires at least two changes. (b) The heat map shows the proportion of inferences where all three ancestral nodes are correctly estimated for the corresponding tree and state pattern. The trees in the test dataset are classified into each of the 8 patterns of internal node states shown in (a) based on their true history. The x-axis shows the tree number, which matches the tree labels in (a), e.g. tree 1 has all internal nodes in state 0. Three methods were used for inference, joint, single node, and marginal. In the joint strategy, a single categorical variable was estimated with possible states 1-8, corresponding the the 8 possible internal node state patterns in (a). For the single node strategy, the node name was given and the ancestral state for that node was estimated as a binary categorical variable. This was repeated three times to estimate the states for all nodes in the tree. In the marginal method, three binary categorical variables were estimated, each corresponding to one of the internal nodes in the tree. For each method, the neural network was trained twice. The numbers on each square match the values shown with the heat map colors.

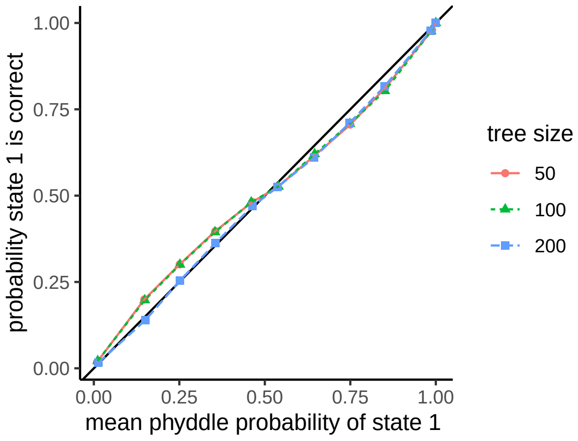


Figure S4: The proportion of nodes where state 1 was the true state by the mean probability for the state reported by PHYDDLE. The lines show the results for three networks trained with different tree sizes. The black line shows $y = x$. If the probabilities were perfectly accurate, the colored lines would fall on the black line.

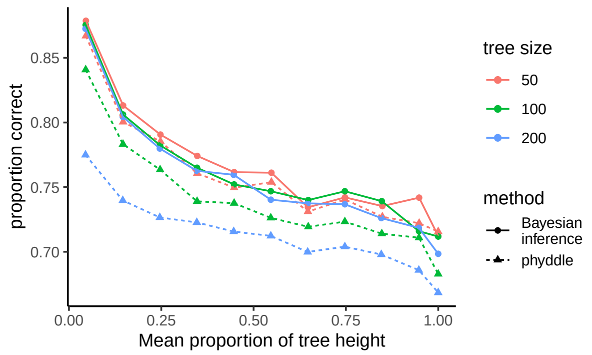


Figure S5: The proportion of inferences where the inferred state is correct by node height relative to tree height for tree with variable sizes. The colors show the maximum tree size and the line and dot types show the method used for inference.

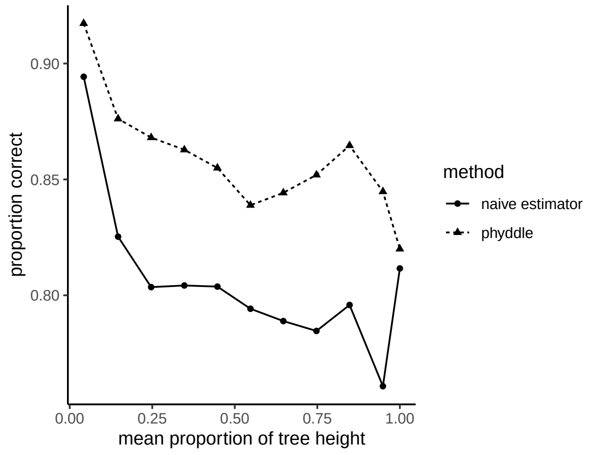


Figure S6: Proportion of inferences that are correct by tree height for PHYDDLE and a naive estimator. The naive estimator infers the ancestral state at a node to be the most frequent tip state among its daughters.

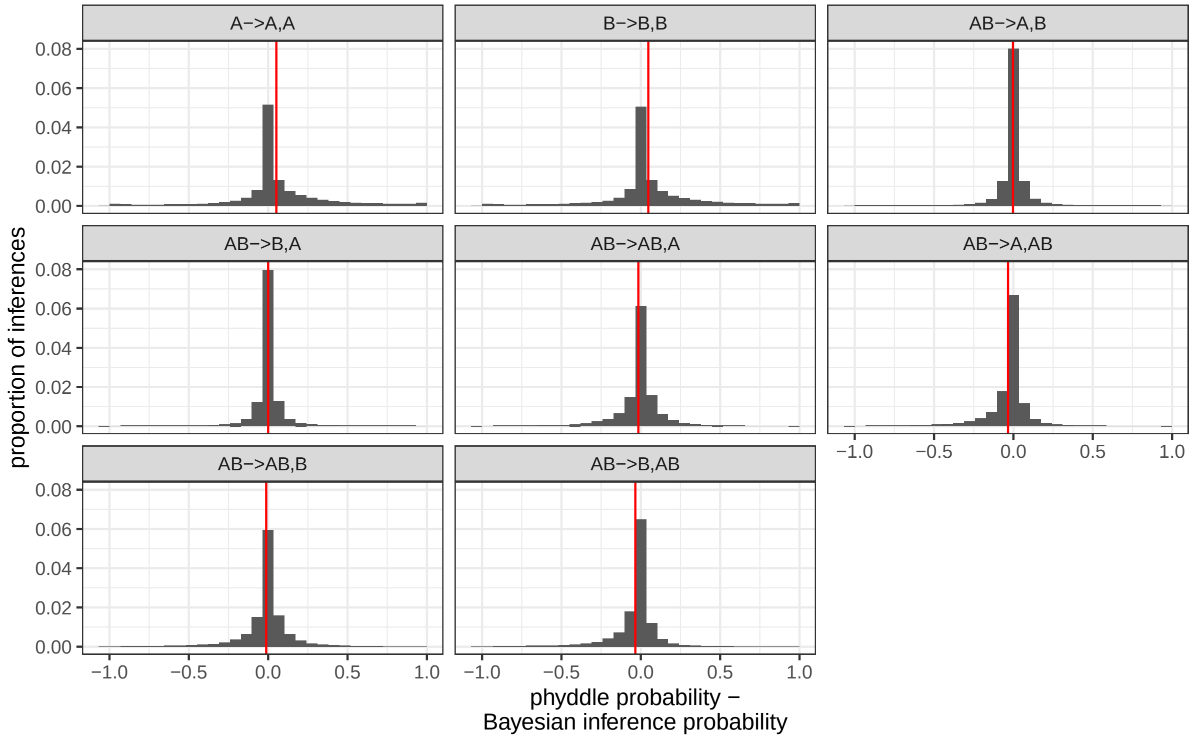


Figure S7: Difference between probabilities reported by PHYDDLE and Bayesian inference for each true state. The red lines show the mean difference for each panel.

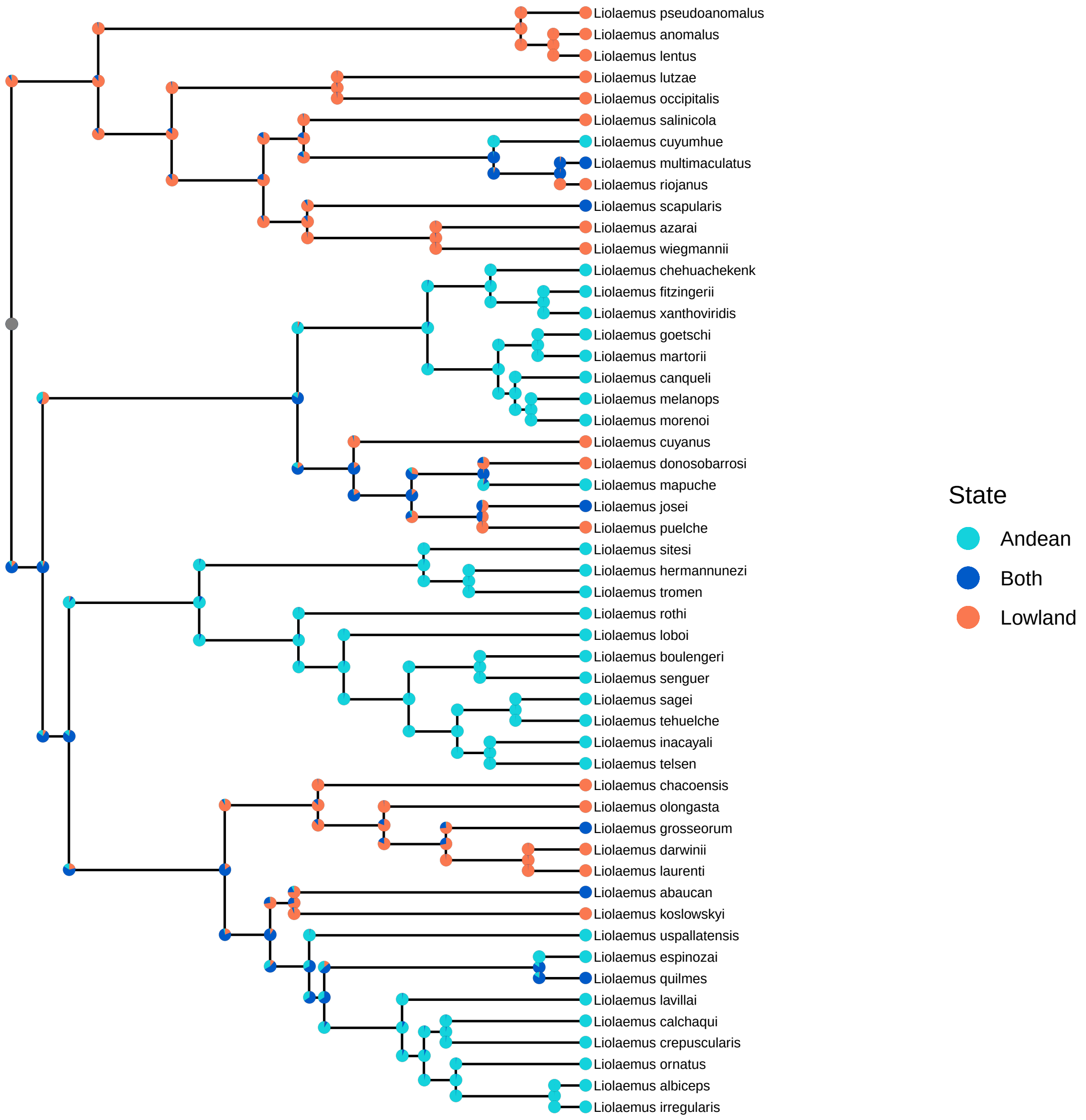


Figure S8: Ancestral state reconstruction for range of *Liolaemus* using RevBayes. The pie charts centered at the internal nodes show the posterior probability of the parent state at the time of speciation. The pie charts on the daughter branches show the posterior probabilities of each of the daughter states immediately after speciation.

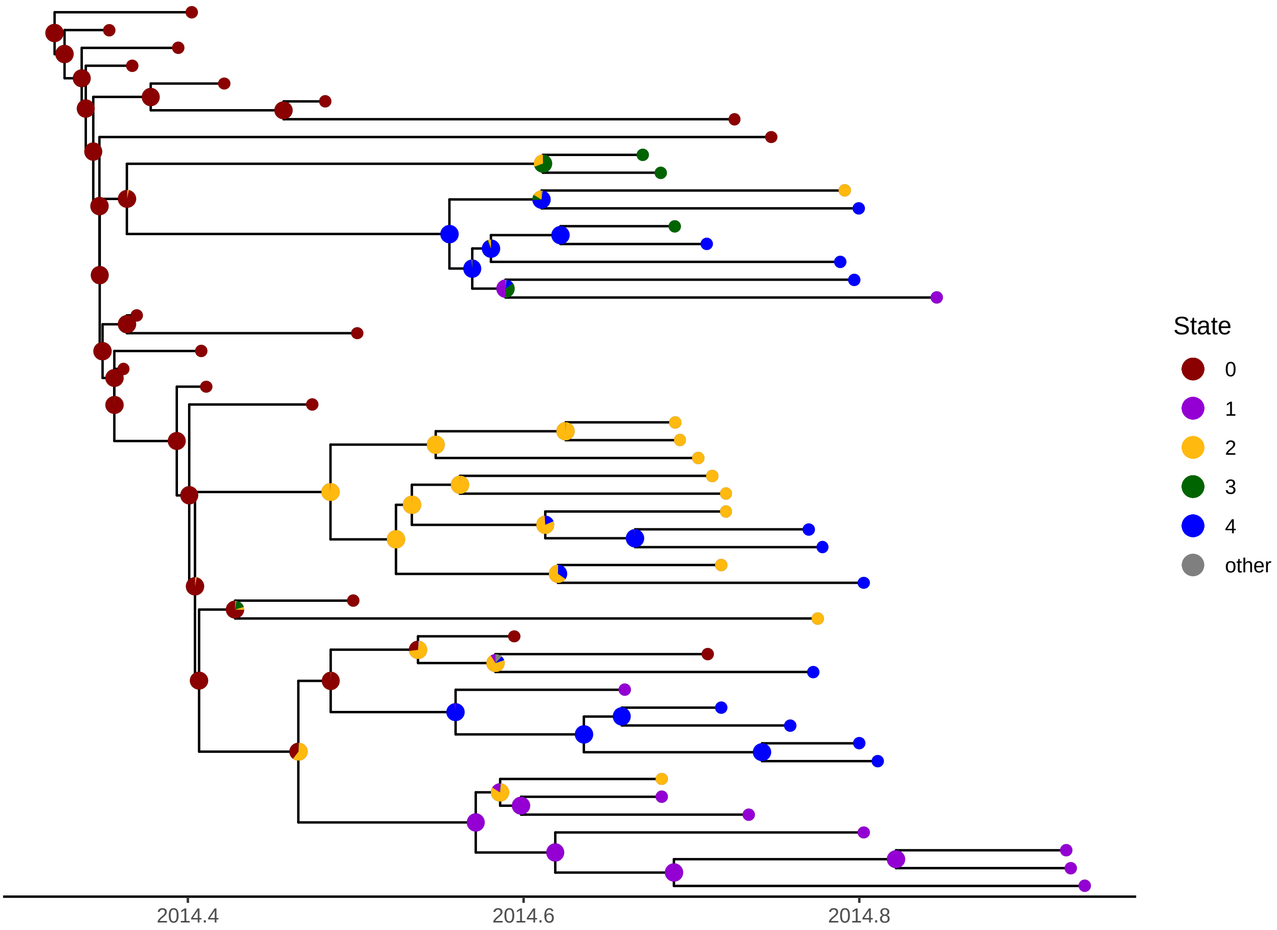


Figure S9: Ancestral state reconstruction of locations from Ebola virus sequences sampled from Sierra Leone during the 2014-2015 outbreak from a first training of the network. The pie charts show the probabilities of each state inferred by PHYDDLE. Locations correspond to groups of districts within Sierra Leon. State 0 corresponds to Kailahun, Kenema, and Bo. State 1 corresponds to Kono, Koinadugu, and Bombali. State 2 corresponds to Tonkolili, Port Loko and Kambia. State 3 corresponds to Pujehun, Moyamba, and Bonthe. State 4 corresponds to Western Rural and Western Urban. Only the three most likely ancestral states are shown by the state color. Any remaining probability for the other two states is shown in gray.

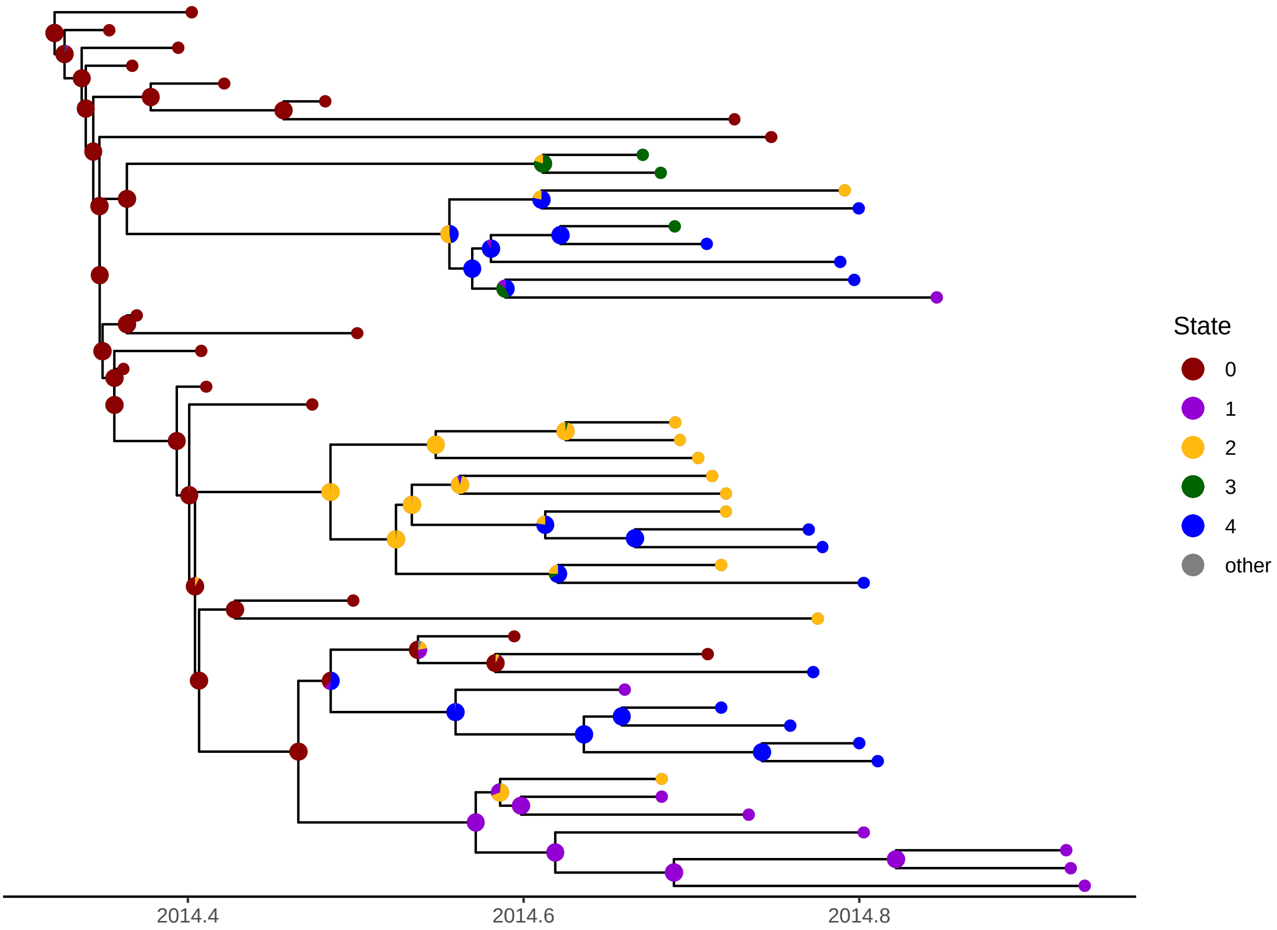


Figure S10: Ancestral state reconstruction of locations from Ebola virus sequences sampled from Sierra Leone during the 2014-2015 outbreak from a second training of the network. The pie charts show the probabilities of each state inferred by PHYDDLE. Locations correspond to groups of districts within Sierra Leon. State 0 corresponds to Kailahun, Kenema, and Bo. State 1 corresponds to Kono, Koinadugu, and Bombali. State 2 corresponds to Tonkolili, Port Loko and Kambia. State 3 corresponds to Pujehun, Moyamba, and Bonthe. State 4 corresponds to Western Rural and Western Urban. Only the three most likely ancestral states are shown by the state color. Any remaining probability for the other two states is shown in gray.

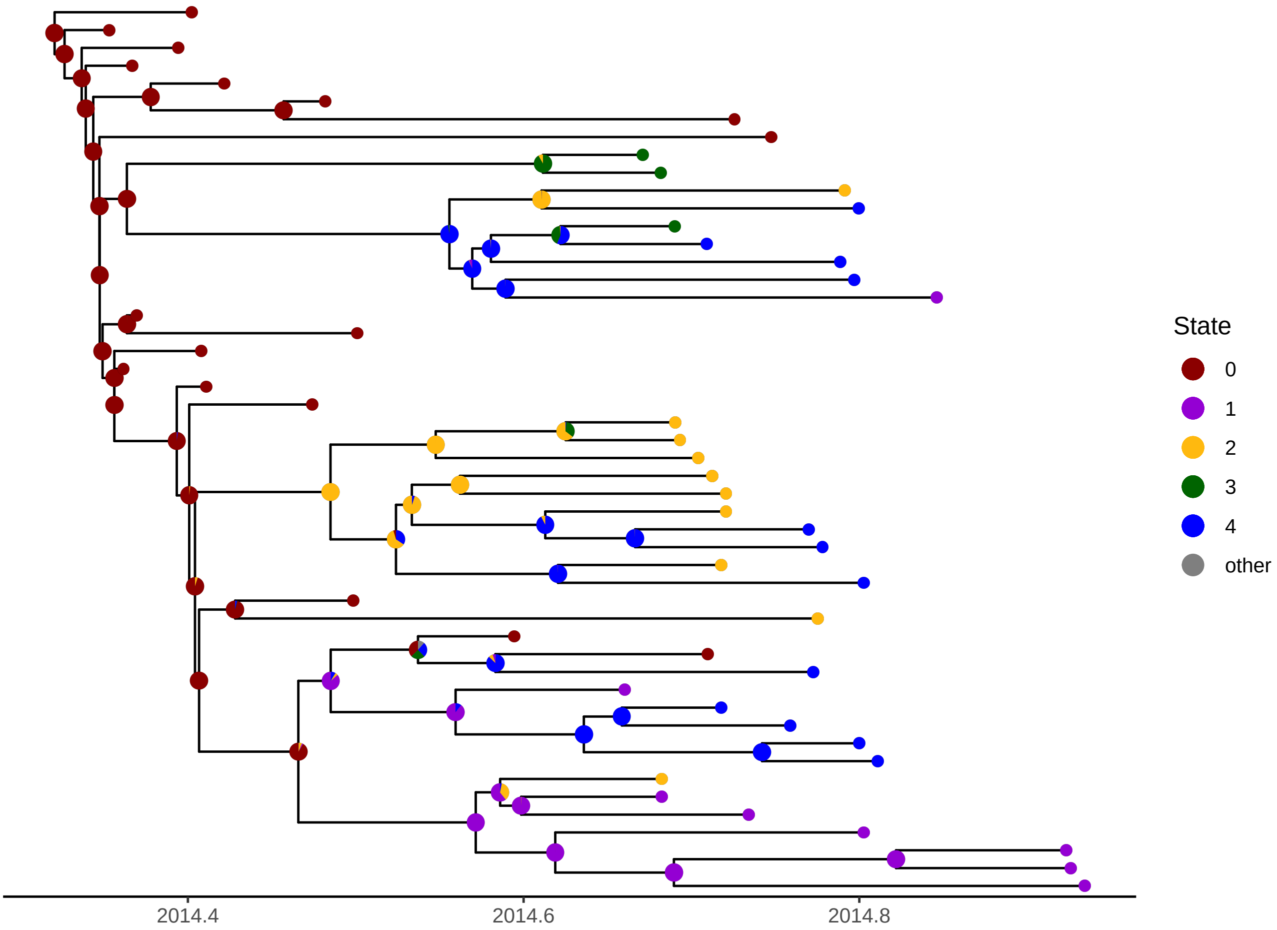


Figure S11: Ancestral state reconstruction of locations from Ebola virus sequences sampled from Sierra Leone during the 2014-2015 outbreak from a third training of the network. The pie charts show the probabilities of each state inferred by PHYDDLE. Locations correspond to groups of districts within Sierra Leon. State 0 corresponds to Kailahun, Kenema, and Bo. State 1 corresponds to Kono, Koinadugu, and Bombali. State 2 corresponds to Tonkolili, Port Loko and Kambia. State 3 corresponds to Pujehun, Moyamba, and Bonthe. State 4 corresponds to Western Rural and Western Urban. Only the three most likely ancestral states are shown by the state color. Any remaining probability for the other two states is shown in gray.

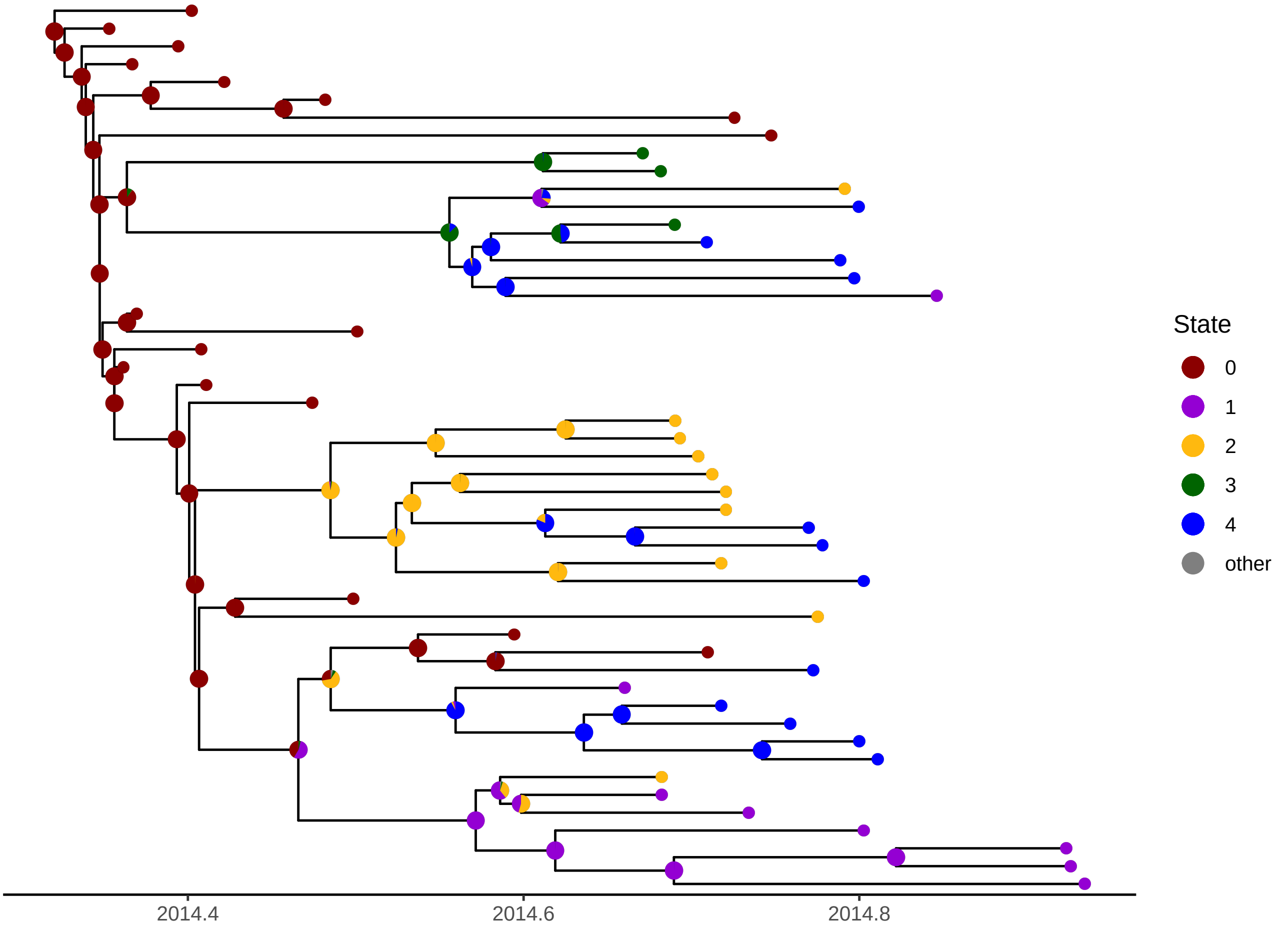


Figure S12: Ancestral state reconstruction of locations from Ebola virus sequences sampled from Sierra Leone during the 2014-2015 outbreak from a fourth training of the network. The pie charts show the probabilities of each state inferred by PHYDDLE. Locations correspond to groups of districts within Sierra Leon. State 0 corresponds to Kailahun, Kenema, and Bo. State 1 corresponds to Kono, Koinadugu, and Bombali. State 2 corresponds to Tonkolili, Port Loko and Kambia. State 3 corresponds to Pujehun, Moyamba, and Bonthe. State 4 corresponds to Western Rural and Western Urban. Only the three most likely ancestral states are shown by the state color. Any remaining probability for the other two states is shown in gray.

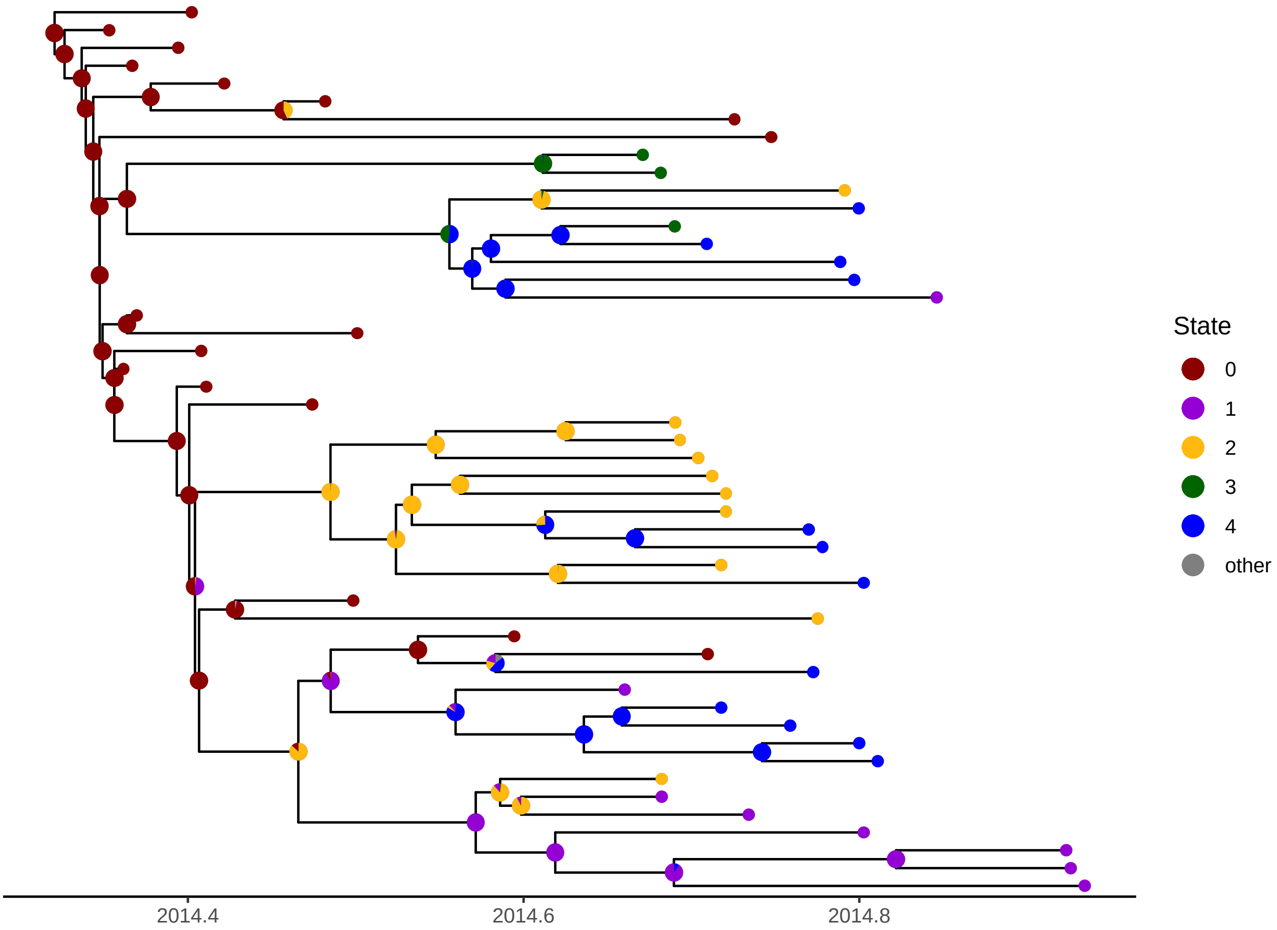


Figure S13: Ancestral state reconstruction of locations from Ebola virus sequences sampled from Sierra Leone during the 2014-2015 outbreak from a fifth training of the network. The pie charts show the probabilities of each state inferred by PHYDDLE. Locations correspond to groups of districts within Sierra Leon. State 0 corresponds to Kailahun, Kenema, and Bo. State 1 corresponds to Kono, Koinadugu, and Bombali. State 2 corresponds to Tonkolili, Port Loko and Kambia. State 3 corresponds to Pujehun, Moyamba, and Bonthe. State 4 corresponds to Western Rural and Western Urban. Only the three most likely ancestral states are shown by the state color. Any remaining probability for the other two states is shown in gray.
